# An embryo-derived peptide signal directs endosperm polarity in Arabidopsis

**DOI:** 10.64898/2025.12.11.693621

**Authors:** Audrey Creff, Jack Rhodes, Camille Salaün, Julien Larive, Vincent Bayle, Emma Turley, Tatsuya Nobori, Duarte D. Figueiredo, Benoit Landrein, Cyril Zipfel, Gwyneth Ingram

**Author notes:** Co-first authors (Audrey Creff, Jack Rhodes).

## Abstract

Angiosperm seed formation requires the coordinated development of the products of double fertilization, the embryo and the endosperm. The endosperm mediates efficient nutrient transfer from surrounding maternal tissues to the developing embryo. This function requires a polarized tissue organization, which manifests as early polar gene expression and polar cellularization dynamics. We show that the receptor kinase HAIKU2 acts in coordination with the transcription factor WRKY10/MINISEED3 to ensure robust endosperm polarity establishment through the activity of the homeodomain transcription factors *WUSCHEL-RELATED HOMEOBOX 8* and *9*.

This process depends on egg cell fertilization and is mediated through the peptide PATHOGEN-INDUCED PEPTIDE-LIKE 7, which acts as a HAIKU2 ligand. Our results reveal how a molecular paracrine dialogue between the embryo and endosperm ensures optimal seed developmental coordination.

## Main Text

In *Arabidopsis thaliana* (hereafter, *Arabidopsis*), seed development is triggered by double fertilization and involves three distinct structures: the embryo (originating from egg cell fertilization), the endosperm (originating from central cell fertilization), and the maternal seed coat. Post fertilization, the endosperm undergoes an initial rapid coenocytic growth phase which, in coordination with the expansion and differentiation of the seed coat, largely determines final seed size (*1–4*). As the embryo reaches the early heart stage, the endosperm cellularizes in a wave-like pattern starting from the distal embryo-surrounding micropylar pole towards the proximal chalazal pole (*5–7*) (Fig 1A). The chalazal endosperm never cellularizes, forming multinucleate cysts of cytoplasm that are proposed to be involved in nutrient uptake from an adjacent proximal vascular unloading zone (*8*, *9*). Endosperm cellularization correlates with seed growth arrest (*10–12*) and the onset of rapid embryo growth, which is supported by a cellularization-associated metabolic shift (*6*, *13*). Thus, endosperm polarity establishment is key to ensure nutrient flow to the embryo. However, although studies in maize have demonstrated a clear influence of the embryo on endosperm development (*14*) to what extent the embryo influences endosperm development in Arabidopsis remains unclear (*15*, *16*).

**Fig 1.**
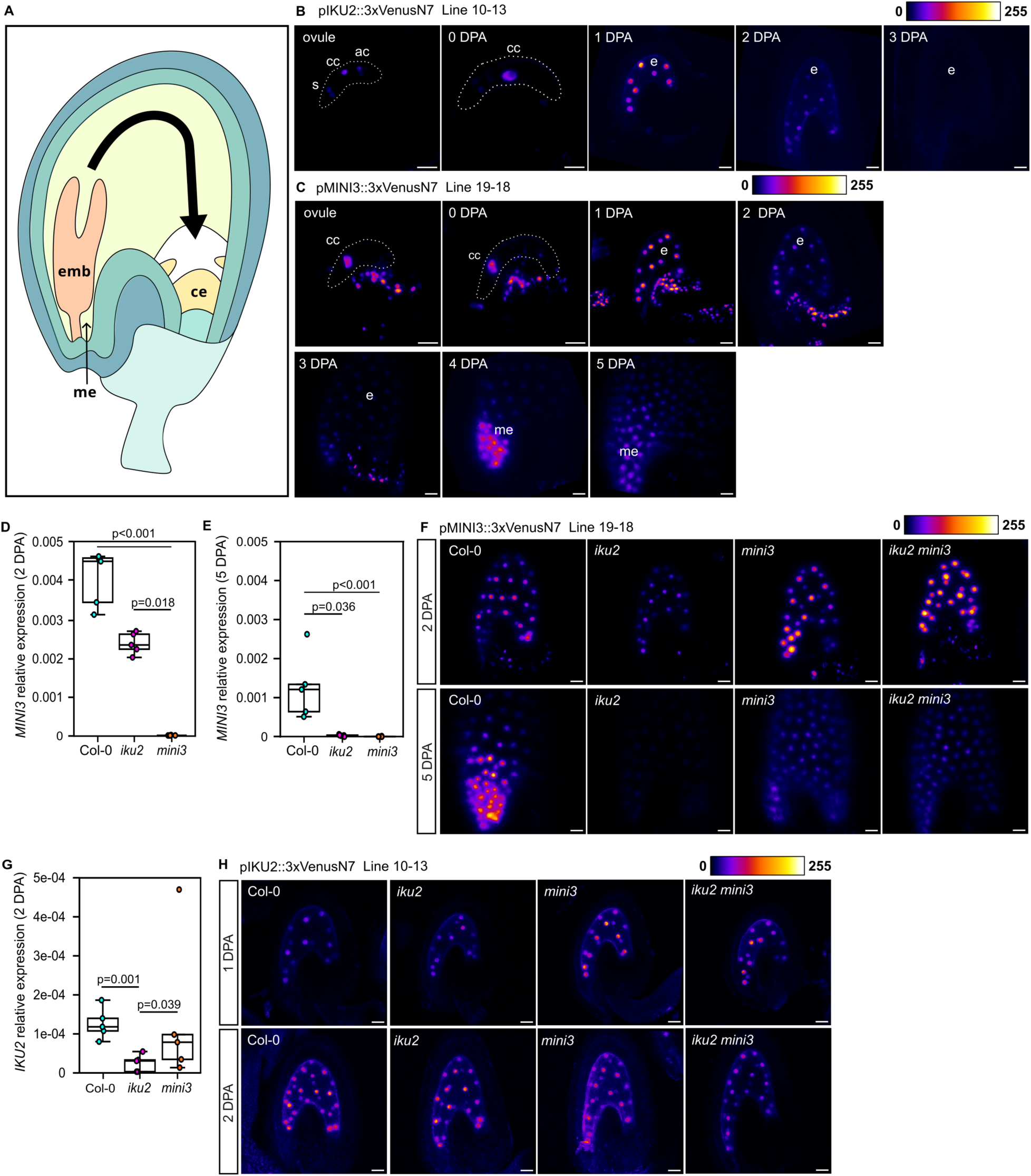
The HAIKU2 RK is required for the polar expression of MINISEED3 in the developing endosperm. (**A**) Cartoon representation of a torpedo-stage Arabidopsis seed showing the main seed tissues. Maternal tissues are shown in shades of blue/green. The embryo (emb), micropylar endosperm (me) and chalazal endosperm (ce) are shown in shades of orange/yellow. The distal-proximal (micropylar-chalazal) polarity of the endosperm is indicated with a filled arrow. (**B**) *IKU2* promoter activity in Col-0 background using a transcriptional reporter line during early seed development. Representative images from n = 6-9 seeds per time point. (**C**) *MINI3* promoter activity in Col-0 background using transcriptional reporter line during early seed development. Representative images from n = 5-8 seeds per time point. (A,B) s, synergid cells; cc, central cell; ac, antipodal cells; e, endosperm; me, micropylar endosperm. Scale bars = 20 µm. (**D**) Relative levels of *MINI3* transcript in 2 days post anthesis (DPA) siliques. (**E**) Relative levels of *MINI3* transcript in 5 DPA siliques. (**F**) *MINI3* promoter activity in single and double mutant backgrounds at 2 and 5 DPA. (**G**) Relative levels of *IKU2* transcript in 2 DPA siliques. (**H**) *IKU2* promoter activity in single and double mutant backgrounds at 1 and 2 DPA. (C,D,F) Transcript levels assessed by RT-qPCR. Each dot represents one plant (n = 5 samples in each case). p-values obtained using the Kruskal-Wallis test followed by Dunn’s Post-hoc test. (E,G) Representative images from n = 10 seeds per genotype per time point. Scale bars = 20 µm.

### HAIKU2 is required for the polar expression of MINISEED3 in the developing endosperm

*HAIKU2 (IKU2)* and *MINISEED3 (MINI3)*, encoding a leucine-rich repeat receptor kinase (LRR-RK) and a WRKY transcription factor (WRKY10) respectively (*11*), play a key role in regulating endosperm cellularization. Loss of either gene causes reduced proliferation and precocious cellularization of the endosperm and reduced seed size). Both *IKU2* and *MINI3* are expressed in the developing endosperm and the two genes are suggested to act in the same pathway (*11*). We found that prior to fertilization, an *IKU2* transcriptional reporter (*3*) is active in the synergids and antipodal cells of the female gametophyte, but not in the egg cell prior to fertilization. After fertilization, expression was observed throughout the endosperm until 2 days post-anthesis (DPA), and then sharply diminished (Figs 1B; S1A). A reporter for *MINI3* expression showed a biphasic expression pattern. Expression was detected in the unfertilized central cell and within some proximal sporophytic cells of the ovule. After fertilization, *MINI3* expression was uniform in the proliferating endosperm until 2 DPA but then diminished drastically, before reappearing specifically around the developing embryo (in the micropylar endosperm) at 4-5 DPA (Figs 1C; S1B). This polar expression is similar to that observed previously (*17*).

To investigate the regulatory relationship between IKU2 and MINI3, we studied *MINI3* expression in *iku2* mutants and *vice-versa*. *MINI3* transcript levels were slightly, but not significantly reduced in *iku2* mutants at 2 DPA (*18*) (Figs 1D; S1C,D). However, transcript levels were significantly lower levels in *iku2* mutant than in wild-type (WT) Col-0 seeds at 5 DPA (Figs 1E; S1F,G). Consistent with these findings, when introduced into the *iku2* background, expression of the *MINI3* reporter at 1-2 DPA was similar to (if slightly weaker than) that in WT seeds, but was strongly reduced compared to WT seeds in the micropylar endosperm at 5 DPA (Figs 1F; S1E,H), suggesting that the second, polar phase of *MINI3* expression depends on IKU2. *IKU2* transcript levels in *mini3* plants at 2 DPA were similar to those in WT seeds (Figs 1G; S1I,J) and were too low for analysis at 5 DPA. When introduced into the *mini3* background, no significant change in expression of the *IKU2* reporter was observed compared to WT seeds (Figs 1H; S1K). Finally, we generated an *iku2 mini3* double mutant and investigated reporter expression in this background. We found that expression of the *MINI3* reporter was similar to that in WT seeds at 2 DPA but almost completely lost at 5 DPA, as observed in *iku2* mutants (Figs 1F; S1E,H). No change in the expression pattern of the *IKU2* reporter was observed in *iku2 mini3* double mutants compared to WT seeds (Figs 1H; S1K).

Our results support a pathway topology in which, contrary to a previous model that placed MINI3 function upstream of the expression of *IKU2* (*11*), *IKU2* and *MINI3* are expressed independently during ovule development and from 1-3 DPA, and both IKU2 and MINI3 are necessary for the establishment of the second, polar phase of *MINI3* expression in the micropylar endosperm from 3-4 DPA.

### IKU2 and MINI3 stabilize endosperm polarity through the activity of WOX8 and WOX9

Consistent with their non-linear regulatory relationship, *iku2 mini3* seed phenotypes indicate both both redundancy and a degree of additivity between *IKU2* and *MINI3* functions. Double mutant seeds were significantly smaller than single mutant seeds (Figs 2A; S2A,E). As previously reported (*11*), the endosperms of *mini3* and *iku2* single mutant seeds almost always cellularized earlier than WT seeds (Figs 2B; S4E). However, in *iku2 mini3* double mutant seeds, endosperm cellularization was heterogeneous, with some seeds cellularizing very early, and others lacking complete cellularization as late as 7 DPA (Figs 2B,D; S4E). Strong cellularization defects correlated with retarded or abnormal embryo development.

**Fig 2.**
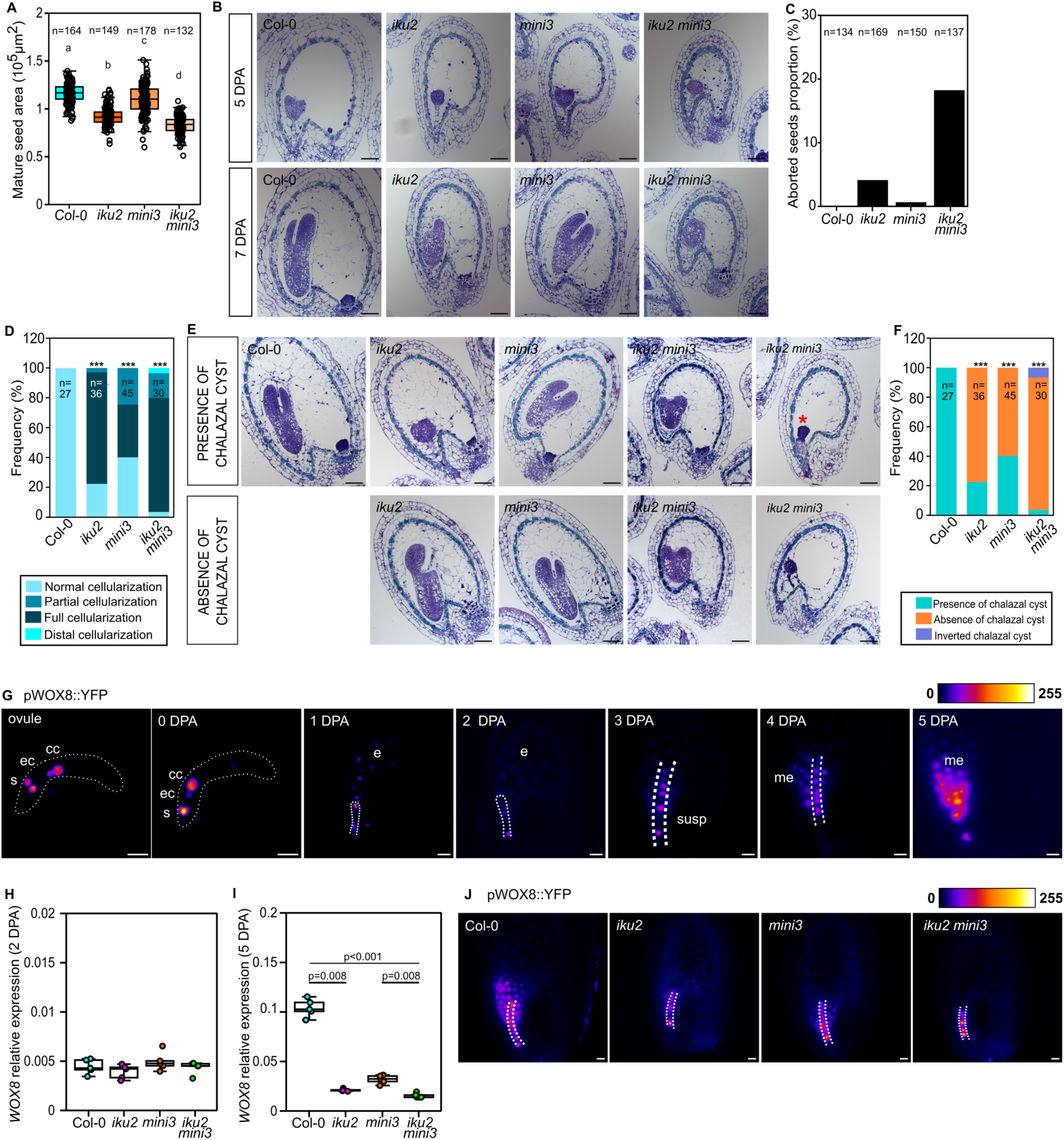
IKU2 and MINI3 stabilize endosperm polarity. (**A**) Measurement of Col-0, *iku2*, *mini3* and *iku2 mini3* mature seed areas from primary inflorescence stems. Total number of seeds is indicated. Statistical groups established using ANOVA with Tukey’s multiple comparison test; p < 0.001. (**B**) Representative images of median sections of Col-0 and mutant seeds stained with toluidine blue at two stages of development. Scale bars = 50 µm. (**C**) Quantification of aborted seeds in Col-0, *iku2*, *mini3* and *iku2 mini3*. The total number of seeds is indicated. (**D**) Patterns of endosperm cellularization in Col-0, *iku2*, *mini3* and *iku2 mini3* in 7 DPA seeds assessed with toluidine blue staining of sections. Data are shown as a contingency bar graph, and the total number of seeds per genotype is indicated. X^2^ test; ^∗∗∗^p < 0.001. (**E**) Representative images of Col-0 and mutant seeds stained with toluidine blue at 7 DPA. Chalazal cyst in micropylar position indicated by a red star. (**F**) Quantification of chalazal cyst presence in Col-0, *iku2, mini3* and *iku2 mini3* seeds obtained from toluidine blue staining of sections. Data are shown as a contingency bar graph, and the total number of seeds per genotype is indicated. X^2^ test; ^∗∗∗^p < 0.001. (**G**) *WOX8* promoter activity in the Col-0 background using a transcriptional reporter line during early seed development; s,synergid cells; cc, central cell; ec,egg cell; e, endosperm; me, micropylar endosperm; susp, suspensor. Representative images from n = 6-8 seeds per time point. (**H**) Relative levels of *WOX8* transcript in 2 DPA siliques. (**I**) Relative levels of *WOX8* transcript in 5 DPA siliques. (H,I) Expression levels were assessed by RT-qPCR. Each dot represents one plant (n = 5 samples in each case). Statistical differences established using the Kruskal-Wallis test followed by Dunn’s Post-hoc test. (**J**) *WOX8* promoter activity in Col-0, *iku2*, *mini3* and *iku2 mini3* at 5 DPA. Embryo suspensors are outlined with dots. Representative images from n = 11-12 seeds per genotype (images shown in Figure S5). (G,J) Scale bars = 20 µm.

Double *iku2 mini3* mutants show high levels of seed abortion (around 20 %), and seed abnormalities (around 65 %) (Figs 2B-F; S2B-D). As reported for single mutants, these phenotypes were dependent upon the zygotic genotype (Fig S3). Levels of abortion and malformation were lower in *iku2* and *mini3* single mutants (Figs 2C,E,F; S2B-D). In addition, the endosperm of both single and double mutants frequently appeared structurally disorganized. Importantly, cellularization of the chalazal endosperm, which is never observed in WT seeds, was complete in the majority (over 75 %) of both developing *iku2 mini3* double mutant and *iku2* single mutant seeds, and almost 40 % of *mini3* mutant seeds (Figs 2D,E; S4) – a phenotype much stronger than that previously reported (*4*). Furthermore, chalazal cysts, which are always present in WT seeds, were predominantly absent in all mutant backgrounds (Figs 2F; S4). Zones of uncellularized (cyst-like) endosperm could occasionally be observed in unexpected positions within the endosperm cavity in some double mutant seeds including around the developing embryo (Figs 2D; S4B). Taken together with the loss of polarized *MINI3* expression, our results suggest that endosperm polarity establishment depends on the HAIKU signalling pathway.

The homeobox transcription factor WUSCHEL RELATED HOMEOBOX 8 (WOX8), acting redundantly with WOX9, has previously been shown to establish embryo polarity (*19*, *20*). *WOX8* is expressed both in the embryo and in the micropylar endosperm (*20*, *21*), and loss of WOX8 function reduces seed size (*21*). *WOX8* expression in the embryo is positively regulated by the transcription factor WRKY2 (*19*, *22*, *23*) which, in turn, is activated via a kinase cascade, likely in response to an unknown endosperm-derived signal (*23–26*). We found that a *WOX8* transcriptional reporter (*19*) was expressed pre-fertilization in the egg cell, synergids and central cell (Fig 2G). After fertilization, expression was observed in the zygote, resolving to the suspensor during early embryo development, as previously reported (*23*). Expression in the endosperm was weak or absent from 1-3 DPA, but increased specifically in the micropylar endosperm at 4-5 DPA (Fig 2G). Although no reduction in *WOX8* transcript levels was observed at 2 DPA in *iku2, mini3*, or *iku2 mini3* compared to WT seeds (Fig 2H), expression was significantly reduced in all mutant backgrounds at 5 DPA (Fig 2I). Consistent with this, although the expression of the *WOX8* transcriptional marker in the embryonic suspensor was maintained in *mini3*, *iku2* and double mutants at 5 DPA, the expression in the surrounding micropylar endosperm became heterogeneous and often very strongly (*iku2 mini3*), strongly (*iku2*) or moderately (*mini3*) reduced in mutants, and was less polar than in WT seeds (Figs 2J; S5). We conclude that IKU2 and MINI3 act together to ensure the robust establishment of polar *WOX8* expression in the micropylar endosperm.

In our growth conditions, we could not detect a strong seed-size phenotype or developmental defects in *wox8* mutant seeds (Fig S6A). However, consistent with publicly available expression data (*27*), *in situ* hybridisation showed *WOX9* expression in the micropylar endosperm, embryo and seed coat (Fig S6B). We measured *WOX9* transcript level in single *iku2*, *mini3* and in double mutants (Fig S6C). At 5 DPA some reduction in *WOX9* transcript levels could be detected in the double mutant compared to WT. *In situ* hybridisation also suggested reduced expression of *WOX9* in the micropylar endosperm of some *iku2* seeds (Fig S6B) indicating that robust polar *WOX9* expression in the micropylar endosperm, like that of *WOX8,* may be promoted by the HAIKU signalling pathway. Expression of *WOX9* in other seed tissues likely masks differences in endosperm transcript levels.

Because single *wox9* mutants show developmental defects and sterility due to both ovule and embryo-developmental defects (*28–30*), and double *wox8 wox9* mutants are embryo lethal (*19*), we assessed endosperm phenotypes of *wox8 wox9* mutant endosperms in homozygous *wox8* mutants heterozygous for *wox9* (*wox8 wox9*/+) at 7 DPA (Figs S7; S8). In seeds with arrested embryos (class B seeds, presumed to contain a *wox8 wox9* mutant embryo and endosperm) the presence of a high proportion of seeds with fully cellularized chalazal endosperms confirmed a redundant role for *WOX8* and *WOX9* in the robust establishment of endosperm polarity. Analyses also revealed likely haploinsufficiency of *WOX9* in the *wox8* mutant background for this phenotype (as observed previously for other phenotypes (*30*)).

Our results show that IKU2 and MINI3 act to establish and stably maintain the expression of *WOX8* and *WOX9* in the micropylar endosperm from 3-4 DPA, and that WOX8 and WOX9 are subsequently required for reproducible differentiation of endosperm tissues.

### IKU2 regulates endosperm polarity in response to a paracrine signal produced by the zygote or the early embryo

Although early Arabidopsis endosperm development was suggested to occur normally in the absence of egg cell fertilization (*15*), a recent study (*16*) identified endosperm-specific genes, including *MINI3,* whose expression depends on the presence of the fertilized embryo. To verify this, we crossed *MINI3* reporter lines with *kokopelli* mutant pollen (*16*, *31*). As previously reported, *kokopelli* crosses led to the production of unfertilized ovules (uo), double fertilized (df) seeds, and seeds in which fertilization had occurred in the egg cell only (eco), or in the central cell only (cco) (Figs 3A; S9A,B,D). At 5 days after pollination (DAP), *MINI3* expression in df seeds was identical to that in WT seeds. No *MINI3* expression was detected in eco seeds. In cco seeds, the expression of *MINI3* was strongly reduced, and when detectable, was distributed uniformly throughout the endosperm (Figs 3A; S9B,C). Similarly, we observed a loss of polar expression of *WOX8* in the endosperm of cco seeds (Fig S9D,E). *IKU2* expression in cco seeds was assessed at 2 DPA, and did not differ from that in double fertilized seeds (Fig S10).

**Fig 3.**
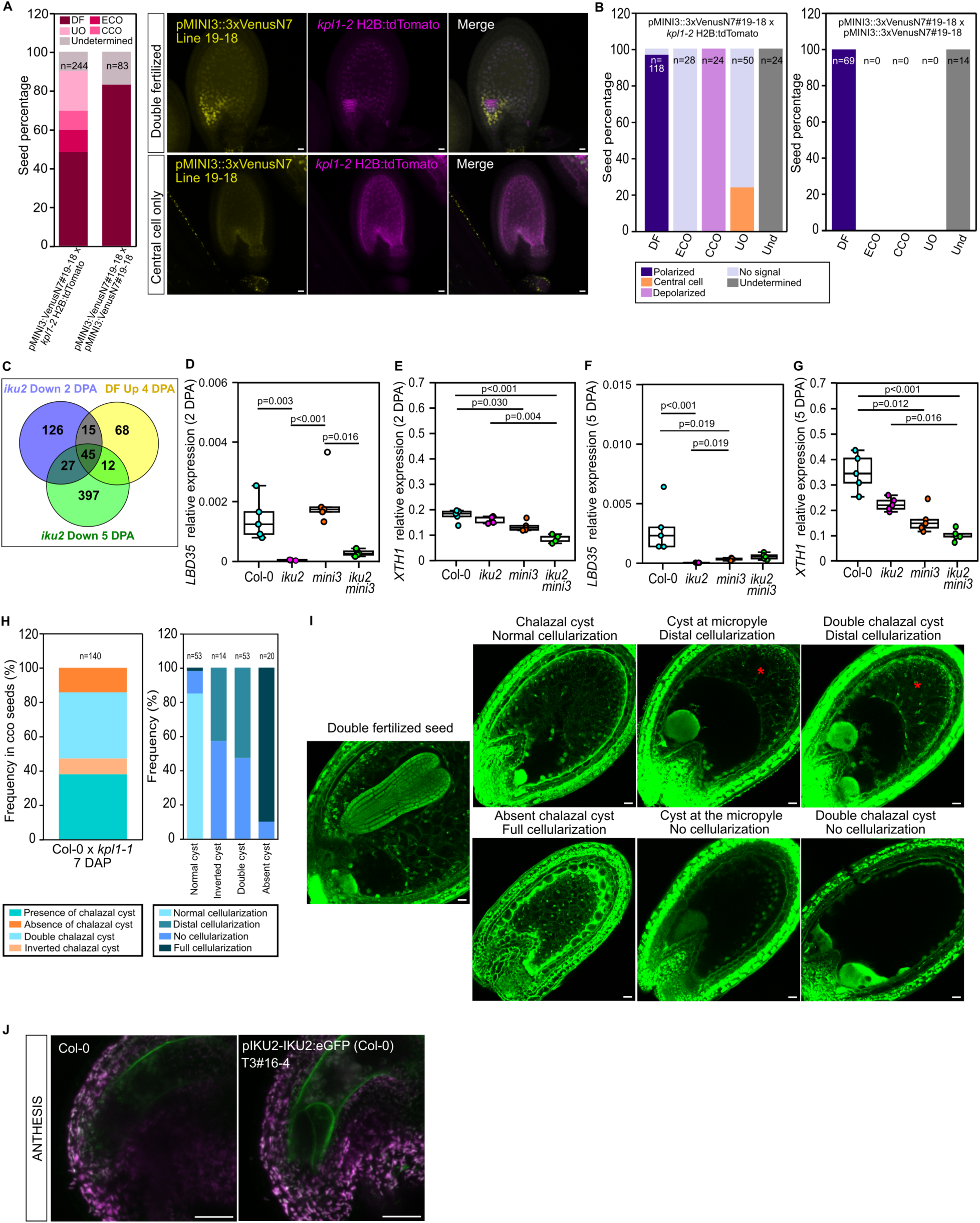
Endosperm polarity depends on egg cell fertilization. (**A**) Left : Quantification of fertilization events from crosses of pMINI3::3xVenusN7 (line 19-18) with *kpl1-2* H2B:tdTomato at 5 DAP. Total number of seeds analysed is indicated. Right: Representative images of *MINI3* promoter activity in double fertilized seeds and cco seeds after Clearsee Alpha treatment at 5 DAP. (**B**) Quantification of signal localization depending upon fertilization events. Total number of seeds analysed is indicated. (**C**) Overlap between genes down-regulated in *iku2* at 2 and 5 DPA and genes up-regulated specifically in double-fertilized seeds at 4 DPA. (*iku2* 2DPA down vs DF up representation factor: 49.9,p < 1.609e-88, *iku2* 5DPA down vs DF up representation factor: 21.0p < 1.697e-60) (**D**) Relative levels of *LBD35* transcript in 2 DPA siliques. (**E**) Relative levels of *XTH1* transcript in 2 DPA siliques. (**F**) Relative levels of *LBD35* transcript in 5 DPA siliques. (**G**) Relative levels of *XTH1* transcript in 5 DPA siliques. (**H**) Left: Presence and position of chalazal cyst within cco seeds from Col-0 x *kpl1-1* cross at 7 days after pollination (DAP). Right: Effect of absence and position of chalazal cyst on endosperm cellularization in cco seeds. Total number of seeds analysed is indicated. (**H**) Representative images of cco seeds harbouring various phenotypes at 7 DPA. Distal cellularization is indicated by a red star. (**J**) IKU2 localisation in wild type plants expressing a *pIKU2-IKU2:eGFP* construct. Representative images at anthesis stage. Scale bars = 20 µm.

RNAseq analysis comparing whole seeds from WT and *iku2* mutant plants was carried out to understand more about the potential function of IKU2-mediated signalling. We identified 252 and 929 differentially expressed genes (DEGs) at 2 DPA and 5 DPA, respectively (Data S1I). DEGs showing reduced expression in *iku2* at 2 DPA were enriched for genes showing specific expression in the early micropylar endosperm (*18*) (Data S1II; Fig S11). These genes also overlapped strongly with genes upregulated only in double-fertilized seeds after *kokopelli* fertilization identified by Zhang and colleagues (*16*) (Figs 3C; S12A, and Data S1III). Furthermore, consistent with IKU2 being necessary for the perception of an embryo derived signal, expression of *LATERAL ORGAN BOUNDARIES 35* (*LBD35*, also called *ASYMMETRIC LEAVES 27, AS27*), previously shown by Zhang and colleagues to be expressed in the endosperm in an egg-cell fertilization-dependent manner (*16*), was fully dependent on IKU2 (Figs 3D,F; S12B), while the expression of *XYLOGLUCAN ENDOTRANSGLUCOSYLASE/HYDROLASE 1* (*XTH1*), which is expressed largely independently of egg cell fertilization, is not strongly influenced by loss of IKU2 (Figs 3E,G; S12C). At 5 DPA, in addition to showing enrichment in micropylar-endosperm expressed genes (including *WOX8* and *WOX9*), DEGs showing reduced expression in *iku2* mutants also contained a second cluster of genes strongly expressed in the chalazal endosperm (Data S1II), consistent with the loss of chalazal cysts in this background.

In summary, transcriptomic analyses suggest that 1) IKU2 is necessary for the expression of genes in the micropylar endosperm, 2) genes showing reduced endosperm expression in the absence of egg cell fertilization also show reduced expression in *iku2* mutants, and 3) IKU2 is necessary for the later expression of genes in the chalazal endosperm.

We next analysed endosperm patterning defects in cco seeds from *kokopelli* crosses. We observed no defects in df seeds. However, consistent with the observed loss of *MINI3* and *WOX8* expression, endosperm development in cco seeds was strongly perturbed (Fig 3H,I). They showed heterogeneity in the timing of cellularization onset, abnormal positioning of zones of uncellularized endosperm within the seed (including complete inversion with a cyst-like zone in the micropylar region, and a fully cellularized chalazal zone), and an apparent reduction in seed size. These phenotypes resemble those observed in developing HAIKU pathway mutants. We also studied the autonomous seeds produced by 50 % of the unfertilized ovules of heterozygous *fertilization independent endosperm (fie)* mutants, which produce endosperms in the absence of central cell fertilization (*32*, *33*). A proportion of these endosperms cellularized (22 % showed cellularization at 8 days after emasculation (DAE)), a phenomenon not previously reported. Abnormal endosperm cellularization phenotypes in *fie* autonomous seeds recapitulated those in cco seeds (Fig S13 A,B).

Our results support the hypothesis that an endosperm-polarizing signal produced by the fertilized egg cell/zygote is perceived by IKU2. We therefore analysed IKU2 localisation in plants expressing a complementing *pIKU2-IKU2:eGFP* construct. GFP signal was observed at the surface of the micropylar central cell/endosperm (Figs 3J; S14), including at the egg cell/zygote - central cell/endosperm interface, at anthesis, consistent with a potential role in the perception of a zygote-derived paracrine signal.

### The PIPL7 peptide is recognized by IKU2 to establish endosperm polarity

IKU2 belongs to LRR-RK subclade XI and forms a monophyletic clade with the closely related paralog RECEPTOR-LIKE KINASE 7 (RLK7), arising from a duplication that likely occurred after the divergence of Ranunculales in the eudicot lineage (*34–38*). RLK7 is the receptor for the family of PATHOGEN-INDUCED PEPTIDE-LIKE (PIPL) peptides, and certain members of the related C-TERMINALLY ENCODED PEPTIDE (CEP) family (*39–43*). To investigate whether PIPL(s) could function as egg cell/zygote-derived signal(s), we interrogated publicly available transcriptomes for *PROPIP(L)* expression. *PROPIPL7* (*AT4G11402*) showed uniquely high expression in zygotes and embryos (Fig S15, S16) (*16*, *44*), especially in zygotes/1-cell embryos after fertilization (*45*). Therefore, *PROPIPL7* became a promising candidate as IKU2 ligand.

Using nuclear-localised transcriptional reporter lines, we showed that *PROPIPL7* expression is detected weakly in synergid cells prior to fertilization (Fig S17) but is strongly and very transiently upregulated in the zygote 1 DPA (Fig 4A, S17). We also observed a very transient and weak signal in endosperm nuclei, and a stronger expression in the seed coat (endothelium) surrounding the micropyle, which was maintained beyond 4 DPA and is consistent with available single nucleus expression data (Fig S17) (*46*).

**Fig 4.**
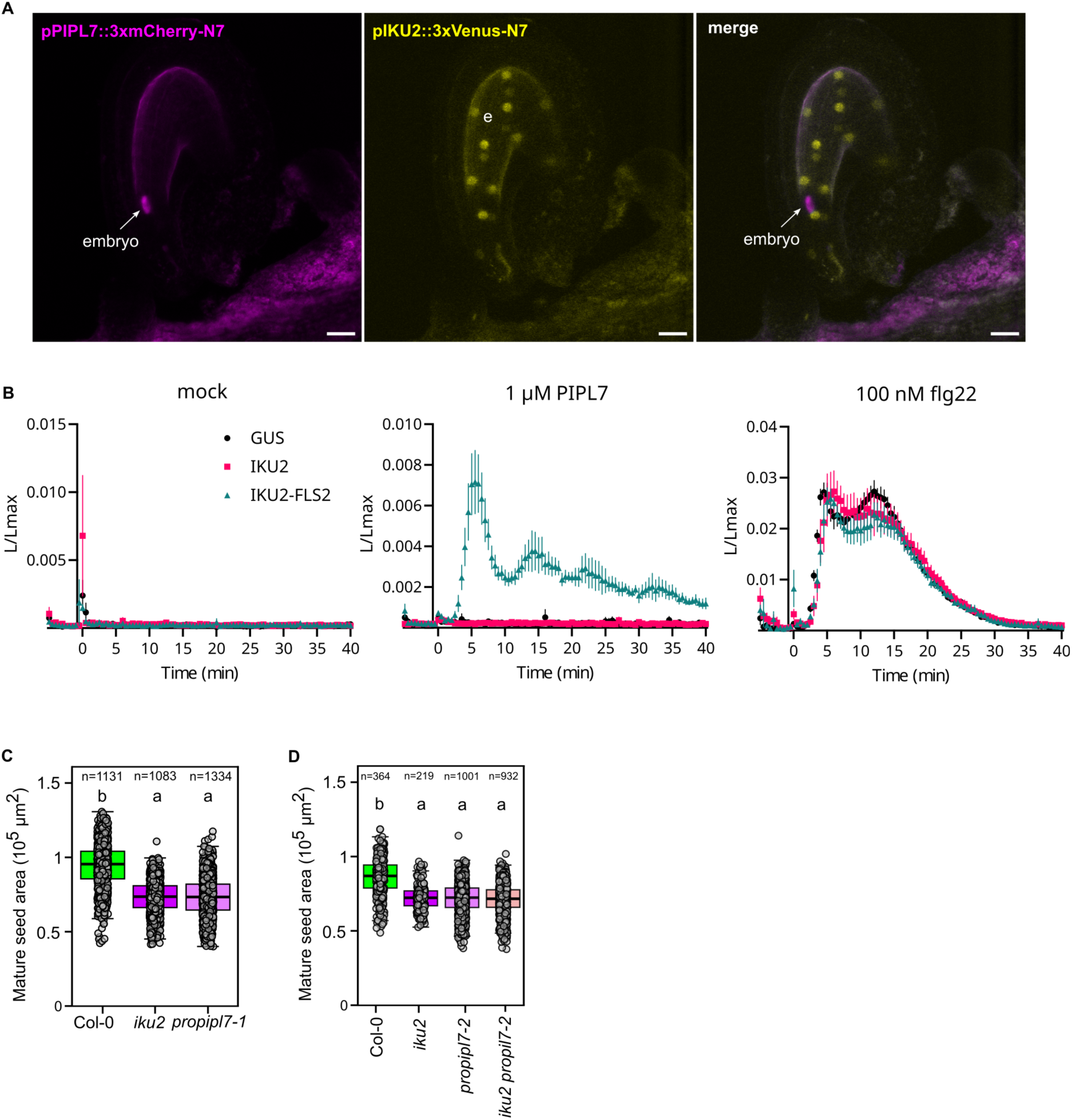
The PIPL7 peptide is recognized by IKU2 to ensure stable endosperm polarity. **(A)** Dual transcriptional reporting of *PIPL*7 (magenta) and *IKU2* (yellow) promoter activity in Col-0 background during early seed development. e, endosperm. Scale bars = 20 µm. **(B)** Receptor-dependent peptide-induced cytoplasmic calcium influx in *N. benthamiana*. IKU2, IKU2-FLS2 or GUS were transiently expressed in 35S::Aequorin transgenic *N. benthamiana* and treated with 1 μM of PIPL7, 100 nM flg22 or mock. Readings were normalised to the maximum value obtained upon discharge. Assays were performed three times with similar results. Error bars indicate S.E.M. n = 4-6 leaf disks. **(C, D)** Measurements of Col-0, *iku2*, *pipl7* and double mutant mature seed areas from primary inflorescence stems. Total number of seeds analysed is indicated. Statistical groups established using ANOVA with Tukey’s multiple comparison test; p < 0.001. Right: Representative images of dry seeds. Scale bars = 200 µm.

To establish whether PIPL7 can function as a ligand for IKU2, we employed a chimeric receptor approach to dissect extracellular perception from downstream signalling outputs (*37*, *47–49*). We fused the ectodomain of IKU2 to the transmembrane and cytoplasmic domain of the LRR-RK FLAGELLIN SENSING 2 (FLS2), which induces potent, ligand (flg22)-dependent, early signalling outputs, such as cytoplasmic calcium influx (*50*, *51*). The chimeric receptor was expressed in a *Nicotiana benthamiana* line encoding a cytoplasmic calcium reporter (Fig S18; (*52*). Whilst PIPL7 treatment did not induce cytoplasmic calcium influx in *N. benthamiana* expressing GUS or IKU2, transient expression of the IKU2-FLS2 chimera resulted in rapid, transient cytoplasmic calcium influx, suggesting receptor-dependent activity (Figs 4B; S19).

Several PIP(L)s were sufficient to induce receptor-dependent responses, suggesting biochemical redundancy within this peptide family (Fig S19), with specific functions dependent rather on unique receptor expression patterns, as shown by *IKU2* in developing seed tissues. We therefore sought to establish genetic evidence that *PIPL7* is linked to IKU2 function, and thus might function as the IKU2 ligand. We generated *propipl7* loss-of-function mutants using CRISPR-Cas9 (Fig S20). Strikingly, *propipl7* mutant seed phenotypes were indistinguishable from those of *iku2* mutants in terms of seed size and development defects, endosperm patterning defects, and alterations in the expression of *MINI3* and *WOX8* (Fig 4C, S21, S22, S23). Phenotypes of *propipl7* mutants, like those of *iku2* mutants, were dependent upon the zygotic genotype (S22). Double *iku2 pipl7* mutants are phenotypically indistinguishable from single mutants (Fig 4C, S21, S23). These results suggest PIPL7 is the IKU2 ligand mediating paracrine signalling between the embryo and the developing endosperm to ensure proper seed development.

Communication between cells and tissues is vital to ensure robust polarisation and development. Double fertilisation results in the fertilisation of the egg cell and central cell by discrete pollen nuclei, which give rise to the embryo and endosperm, respectively. These tissues must coordinate development to ensure successful reproduction. Over two decades ago, forward genetic screens identified mutants in the HAIKU signalling pathway, which impair seed development. Here, through detailed characterisation of the pathway and synthetic biology approaches, we identify the upstream signal as PIPL7, a peptide derived from the fertilised egg-cell lineage, which is required for reproducible endosperm polarisation and cellularisation. Our findings therefore provide novel insights into the molecular mechanisms underlying the appropriate development of seeds – which are the source of most of our food. Furthermore, by defining one of the last remaining orphan receptors from the LRR-RK subfamily XI, this study opens the door for a better understanding of the co-evolution of plant ligand-receptor specificity and function.

## Acknowledgments

The authors thank Frederic Berger (Gregor Mendel Institute of Molecular Plant Biology,Austria), Thomas Laux (University of Freiburg, Germany), the Stegmann lab (Ulm University, Germany), Ryoshiro Kasahara (Nagoya University, Japan) and Masaru Ohme-Takagi (Saitama University, Japan) for kindly sharing genetic materials, and Jeremy Just for help with transcriptomic data analysis. We that Sara Simonini and Ueli Grossniklaus for helpful comments on the manuscript. We acknowledge the contribution of SFR Biosciences (Université Claude Bernard Lyon 1, CNRS UAR3444, Inserm US8, ENS de Lyon) PLATIM-LyMIC for technical assistance with microscopy. We thank the RDP Plant Culture team (Alexis Lacroix, Patrice Bolland, Camille Knaupp, Joseph Pacitto and Justin Berger) and the John Innes Centre Horticultural Services (especially T Wells; J Taylor, K. Bachowska) for plant care and A Wawryk from the TSL Plant Transformation support group for plant transformation. We thank Isabelle Desbouchages and Hervé Leyral for technical assistance regarding molecular biology work; and Cindy Vial, Laureen Grangier, Nelly Camilleri, Stéphanie Maurin and Julie Prata for administrative assistance. We thank all past and current members of the Zipfel and Ingram groups for technical help and fruitful discussions.

## Funding

Ecole Normale Supérieure de Lyon (GI, BL) Gatsby Charitable Foundation (CZ) University of Zurich (CZ)

BBSRC Institute Strategic Programme Grant in Advancing Plant Health (BB/Y002997/1) (CZ)

The European Research Council under the European Union’s Horizon 2020 research and innovation programme no. 773153 (project ‘IMMUNO-PEPTALK’) (CZ), Max Planck Society (DF)

German Federal Ministry for Education and Research grant number 031B1230A (DF)

## Author contributions

Conceptualization: GI, CZ, JR, TN Design of the work; GI, AC, CZ, JR,

Acquisition of data; AC, JR, ET, CS, JL, BL Analysis of data; AC, JR, CS, BL Interpretation of data; GI, CZ, AC, JR, BL, DF Creation of new software used in the work: VB Writing: GI, AC, CZ, JR, BL

## Competing interests

Authors declare that they have no competing interests.

## Data and materials availability

All data are available in the main text or the supplementary materials.

## Supplementary Materials

Materials and Methods Supplementary Text

Figs S1 to S23 Table S1

Data S1 to S3

## Notes

### Competing Interest Statement

The authors have declared no competing interest.

