## supplementary materials for "An embryo-derived peptide signal directs endosperm polarity in Arabidopsis"

**The PDF file includes:**

Materials and Methods
Figs. S1 to S23
Table S1

**Other Supplementary Materials for this manuscript include the following:**

Data S1 to S3

#### Materials and Methods

##### Materials

Plant Materials

The Columbia-0 (Col-0) *Arabidopsis thaliana* accession was used in this study. A list of all the mutant lines used are provided in the key resources table and were obtained from the Arabidopsis Biological Resource Center (ABRC; Ohio, USA) (*mini3-2* (*SALK_050364*) or directly from Frédéric Berger (*iku2-2* (*11*, *53*) introgressed into Col-0) (Gregor Mendel Institute of Molecular Plant Biology, Austria), Thomas Laux (*wox8-3*, *wox9-1* (*19*) and *proWOX8::YFP)*(University of Freiburg, Germany), and Ryoshiro Kasahara (Nagoya University, Japan) and Masaru Ohme-Takagi (Saitama University, Japan) (*RPS5Apro::H2B:tdTomato/kpl*). *iku2 mini3* double mutants were generated by crossing *iku2-2* and *mini3-2* (SALK_050364). The *fie-12* allele used in this study is described in (*54*). Homozygous mutants were identified by genotyping with primers in Table S1. Fluorescent lines used for this study include *pIKU2::3X-Venus-N7* (*3*), *RPS5Apro::H2B:tdTomato/kpl* (*16*, *55*), and *proWOX8::YFP* (*19*). All other reporter lines were developed specifically for this study (see below).

Growth conditions and seeds staging

Seeds were gas sterilized in open Eppendorf tubes with chlorine (generated by mixing 4 mL HCl (37 %) in 100 mL bleach in a closed container) for 2 h and sown on plates with Murashige and Skoog (MS) medium and 0.5 % sucrose in sterile conditions, stratified for 2 days at 4 °C and grown for 11 days in a Sanyo (Fisher Scientific) growth cabinet under short-day conditions (8 h light, 21 °C, 150 $\mu mol.m^{-2}.s^{-1}$ during the day, 18 °C during the night). Seedlings were then transferred into separate pots of soil (Argile 10 (Favorit)), and placed in a short-day growth chamber (8 h light, 21 °C and 150 $\mu mol.m^{-2}.s^{-1}$ during the day, 18 °C during the night) for 10 days before being transferred to a long day growth chamber (16 h light, 21 °C, 150 $\mu mol.m^{-2}.s^{-1}$) to induce flowering. Seeds were staged every day by marking the newly opened flowers with coloured cotton threads.

##### Methods

Genotyping

Plant genomic DNA was extracted from rosette leaves with a rapid CTAB isolation technique as described by Stewart and Via (*56*). Leaves were first ground with liquid nitrogen and resuspended in 300 µL of extraction buffer (CTAB 2 %, NaCl 1.4 M, EDTA pH8 20 mM, Tris-HCl pH 8 100 mM; all purchased from Sigma) and incubated for 20 min at 60 °C. After addition of 300 µL chloroform (VWR) and centrifugation for 20 min at 4000 rpm, gDNA was precipitated from the supernatant using 2-propanol (Merck) and pelleted. The pellet was washed with 70 % ethanol and dried for 15 min at 60 °C. The pellet was resuspended in 100 µL of TE with 10 µg/mL of RNase A and incubated for 30 min at 37 °C. DNA (1 µl) was used to perform PCR reactions using GoTaq G2 (Promega). *iku2-2*, *mini3-2*, *wox8-3* and *wox9-1* and *fie-12* mutant alleles were genotyped with primers listed in Table S1.

RNA extraction and cDNA synthesis for RT-qPCR experiments

Total RNA from siliques at 2 and 5 DPA was extracted using the Spectrum Plant Total RNA Kit (Sigma-Aldrich) according to the manufacturer's instructions, and eluted in a 50 µL RNase-free elution buffer. Total RNAs were digested with Turbo DNA-free DNase I (Invitrogen) according to the manufacturer's instructions. The mRNAs were reverse transcribed using the SuperScript VILO cDNA Synthesis Kit (Invitrogen) according to the manufacturer's protocol and freshly synthesised cDNA was diluted by a factor of 1:20 in milliQ $H_2O$ prior to qPCR analysis.

Quantitative gene expression analysis

PCR reactions were performed in an optical 384-well plate in the QuantStudio™ 6 Flex Real-Time PCR System (ThermoFisher Scientific), using FastStart Universal SYBR Green Master (Rox) (Roche), in a final volume of 10 µL, according to the manufacturer's instructions. The following standard thermal profile was used for all PCR reactions: 95 °C for 10 min, 40 cycles of 95 °C for 10 s, and 60 °C for 30 s. The data were analysed using the QuantStudio Real-Time PCR Software v1.3 (Applied Biosystems). As a reference, a geometric mean between two house-keeping genes, *EIF4A1* and *AP2M*, was used to normalise gene expression. For each primer pairs, PCR efficiency (E) was estimated from the data obtained from standard curve amplification using the equation $E = 10^{-1/\text{slope}}$. Expression levels are presented as $E^{-\Delta Ct}$, where $\Delta Ct = Ct_{GOI} - Ct_{REF}$. The sequences of the primers used for qPCR can be found in Table S1. Each experiment contains 5 samples per genotype per condition and was repeated three times with independent biological samples (i.e., from independent batches of plants growing at different times).

Generation of RNAseq data

Total RNA from developing seeds at 2 and 5 DPA was extracted and treated as described in the previous section. Three independent biological replicates were produced. The following steps were carried out by the HELIXIO sequencing company (Biopôle Clermont-Limagne, Saint-Beauzire, France). RNA purity was checked using a spectrophotometer NanoDrop ND-1000 (Thermo Fisher Scientific), RNA quantity was measured using a Qubit® 2.0 fluorometer (Thermo Fisher Scientific), and RNA integrity was validated thanks to an analysis of the ribosomal RNA using a Bioanalyzer 2100 (Agilent Technologies). The cDNA libraries were then produced using the following kits:  NEBNext Ultra II Directional RNA Library Prep Kit for Illumina (New England Biolabs), NEBNext Poly(A) mRNA Magnetic Isolation Module » (New England Biolabs) and NEBNext Multiplex Oligos for Illumina (New England Biolabs). Single read sequencing (75bp) was performed using a NextSeq500 (Illumina). The reads were aligned against the reference genome (Arabidopsis thaliana TAIR 10, release 42) and quantified using the STAR program (https://github.com/alexdobin/STAR) and the cufflink method (http://cole-trapnell-lab.github.io/cufflinks). Analysis of the differentially expressed genes was done using the DeSeq2 library in R. The overlap between different transcriptomic datasets were visualised using the Venny software (https://bioinfogp.cnb.csic.es/tools/venny/). The measurements of the statistical significance of the overlap between two groups of genes were done using the following software: http://nemates.org/MA/progs/overlap_stats.html, using a total number of *Arabidopsis* genes of 24'797 (based on the number of genes found in our transcriptome). The analysis of the spatial expression of the genes based on available microarray data (*27*) was done using the eNorthern expression browser of BAR (https://bar.utoronto.ca/affydb/cgi-bin/affy_db_exprss_browser_in.cgi). A reclustering of the data based on Euclidian distance and heatmap visualization was then performed in R. Analysed datasets can be found in Data S1.

Cloning and transgenic line generation

All primers and plasmids are listed in Table S1. The *MINI3* promoter was amplified using Q5 High-fidelity DNA polymerase (New England Biolabs) with the primers Prom-MINI3-B4 and Prom-MINI3-B1R and cloned into *pDONR-P4-P1R* (Life Technologies). A triple LR Gateway reaction (Life Technologies) was then performed using the *pMINI3-pENTR-R4-L1*, *3X-VENUS-N7-pENTR-L1-L2*, and *3'-ter-pENTR-R2-L3* plasmids as entry vectors and the *pH7m34GW* plasmid (*57*) as a destination vector to generate a *pMINI3::3X-VENUS-N7-pH7m34GW* construct (conferring Hygromycin resistance in plants). Plasmids were amplified and purified from subcloning efficiency TOP 10 *E. coli* cells (Invitrogen) using the NucleoSpin Plasmid Kit (Macherey-Nagel) according to manufacturer's instructions. The final destination construct was transformed into *Agrobacterium* strain GV3101 (GoldBio) via electroporation using the floral dip method (*58*) .

The 1245-bp promoter of *PROPIPL7* was synthesized by Genewiz GmbH and cloned into the vector pUC-GW-Amp flanked by BsaI sites (CZLp7320), which allowed release of the promoter sequence flanked by GGAG-AATG overhangs for integration into Goldengate modular cloning plans (*59*). This fragment was integrated into the preassembled backbone (CZLp7748) which contains a 3xVenus-N7 CDS (*60*) and HSP18.2 (pICSL60028) terminator alongside a kanamycin selection cassette. This vector was used to transform wild-type Arabidopsis (Col-0).

The 3xmScarlet3-N7 coding sequence (CZLp8275) was assembled from PCR-amplified fragments into the acceptor plasmid pICH41308. Together with the previously described 3xmCherry-N7 reporter (*61*), these fluorescent reporter modules were assembled into transcription units driven by the *PROPIPL7* promoter (CZLp7748) and terminated by the HSP18.2 terminator (pICSL60028). The module was assembled in the pICSL86900 backbone, which carries a kanamycin-resistance cassette for selection in transgenic plants. These constructs were subsequently introduced into the preexisting *pIKU2::3xVENUS-N7* line (*3*).

The *IKU2* sequence was amplified from genomic DNA with the indicated primers to allow the domestication of BsaI sites and to allow direct cloning into the expression vector pICSL86977 with the pICSL50044 mEGFP C-terminal tag to generate p35S::IKU2-mEGFP (CZLp4332). The IKU2-FLS2 chimera was cloned using a similar approach with PCR amplification generating products for a scarless BsaI restriction-ligation reaction into pICSL86977, using an alternative GFP variant, pICSL50064 (mCitrine), as the C-terminal tag.

To mutate *PROPIPL7* gRNAs were designed using CHOPCHOP to target the *PROPIPL7* coding sequence (*62*). gRNAs were assembled into transcriptional units using pU6 promoters Goldengate cloning following the approach described by Castel et al. (*63*). Concatenated gRNAs were then assembled into a final LP expression vector (CZLp6841) with a polyintronic Cas9 nuclease driven by the RPS5a promoter (*64*) and FastRed seed selection marker derived from pICSL33004. Mutants were confirmed using Sanger sequencing (Genewiz GmbH). As the CRISPR construct also targeted the *PROCEP16/PROPIPL1/ AT1G49800* CDS the single *propipl7* mutant was isolated through backcrossing to the Col-0 wild-type to segregate away *propipl1* mutations and selected to homozygosity. Sequences of the alleles can be found in Data S2 and Data S3.

<u>Seed clearing and size measurements</u>

At 8 DPA, siliques were opened with a needle, and the seeds were removed with forceps and put in a drop of clearing solution (1 vol glycerol/7 vol chloral hydrate liquid solution, VWR Chemicals) between a slide and a coverslip. The samples were incubated for at least 24 h at 4 °C before being imaged with a Zeiss Axioimager 2 equipped with a 20× DIC dry objective. The resulting images were semi-automatically analysed using a specifically developed macro (https://github.com/RDP-vbayle/SiCE_FIJI_Macro/blob/main/VariousSeedDev/MacroClearing-Creff_et_al_2024.ijm) in the Fiji (*65*) software. For each segmented seed, the embryo phenotype was scored, and seed area, major and minor axes were measured. For mature seeds measurements, the seeds were imaged using a Leica stereomicroscope. The resulting images were automatically analysed using a specifically developed macro in the Fiji software(https://github.com/RDP-vbayle/SiCE_FIJI_Macro/blob/main/VariousSeedDev/MacroClearingLeica-Creff_et_al_2024.ijm). Seeds that were too close to each other to be segmented separately were manually removed from the analysis after segmentation.

<u>Adapted ClearSee Alpha protocol for siliques</u>

ClearSee solution was prepared as described in (*66*, *67*) by mixing xylitol powder 10 % (w/v) final concentration (Sigma-Aldrich), sodium deoxycholate 15 % (w/v) final concentration (Sigma-Aldrich) and urea 25 % (w/v) final concentration (Euromedex) in water. Sodium sulfite (0.73 %) (Sigma-Aldrich), was added to the ClearSee solution just before each use (ClearSee Alpha). Siliques were opened with a needle along the septum and fixed in 4 % (w/v) paraformaldehyde (Sigma-Aldrich) in 1X PBS buffer pH 6.9 under vacuum (700 mmHg) for 1 h on ice, breaking and replacing the vacuum at once after 30 min. The fixative was replaced with a fresh aliquot, and samples were incubated overnight at 4 °C. Fixed tissues were washed twice for 1 min in 1X PBS and cleared with ClearSee Alpha at room temperature for two weeks protected from light under gentle agitation. The solution was changed with fresh ClearSee Alpha every two days.

<u>LR (London Resin) white embedding, sectioning and Toluidine blue staining</u>

Seeds were fixed in ice-cold PEM buffer containing 50 mM PIPES (Sigma-Aldrich), 5 mM EGTA (Sigma-Aldrich) and 5 mM $MgSO_4$, pH 6.9 (Sigma-Aldrich) with 4 % (w/v) paraformaldehyde. The samples were placed under vacuum (2 × 30 min on ice) followed by overnight incubation at 4 °C in fresh fixative solution. After rinsing twice in PEM buffer, samples were dehydrated through an ethanol series (50 %, 70 %, 80 %, 90 % and 100 % three times) under vacuum and infiltrated with increasing concentrations (30 %, 50 % and 100 %) of LR White resin (London Resin Company) in absolute ethanol over 8 days before being polymerized at 60 °C for 24 h. The samples were sectioned (1.0 µm thickness) using a diamond knife at a 45° angle (Diatome, LFG Distribution) mounted on a Leica RM2265 microtome, and dried onto glass slides at 70 °C.

For toluidine blue staining, the sections were incubated for 20 s at 70 °C with filtered Toluidine blue 1 %/borax 1 % before being rinsed with distilled water, dried and mounted in Entellan mounting medium (Merck). The sections were imaged with a Zeiss Axioimager 2 equipped with a 20x dry objective.

<u>Feulgen staining</u>

The siliques were opened and fixed overnight at 4 °C in a 3:1 mixture of absolute ethanol:glacial acetic acid. The fixative was replaced with 70 % ethanol and the siliques were stored at 4 °C until needed. The samples were rinsed three times with distilled water for 15 min and then hydrolysed in 5 N HCl for 1 h at room temperature. After hydrolysis, the siliques were rinsed three times with distilled water for 15 min and then stained with Schiff's reagent (Sigma-Aldrich) for 3-4 h at room temperature. Samples were then rinsed three times for 15 min with cold distilled water. The samples were then dehydrated in ethanol baths of increasing concentration (30 %, 50 %, 70 %, 96 %) for 10 min each at room temperature and then stored in absolute ethanol overnight at 4 °C. Absolute ethanol baths were repeated until the bath remained colourless. The samples were brought to room temperature, and the ethanol was replaced with a 70: 30 mixture of absolute ethanol: LR White (Electron Microscopy Sciences) for 1 h. The bath was replaced by a 50:50 then 30:70 mixture of absolute ethanol:LR White at room temperature for 1 h. The samples were then placed in pure LR White at room temperature. The bath was renewed after 1 h and the samples were placed at 4 °C overnight. The siliques were dissected, and seeds were placed between slide and coverslip in LR White containing the polymerisation catalyst from the kit (9.9 g benzoyl peroxide) for polymerisation. The slides were incubated at 60 °C for 16 h. The coverslips were then removed with a razor blade, and the slides were kept in the freezer until they were observed under a confocal microscope.

Confocal microscopy

For promoter activity, individual seeds were placed on adhesive tape on a microscope slide and covered with water. Z-stacks of Col-0 and mutant seeds expressing *pIKU2::3X-Venus-N7*, *pMINI3::3xVenusN7* and *pWOX8::YFP and pPROPIPL7::3xVenus-N7* were acquired using a Leica SP8 upright confocal microscope equipped with a 40x water immersion objective (HCX APO L UV 40 x 0.8 W wd 3.3 mm Deeping (Leica #11506155) or a 25x water immersion objective (HCX Fluotar VISIR 25 x 0.95 W wd 2,5mm Deeping (Leica#11506507). Venus and YFP were excited with an LED laser emitting at a wavelength of 514 nm (Leica Microsystems). The signal was collected at 520–550 nm.

The following scanning settings were used: pinhole size 1AE, 1.25× zoom, 20 % laser power, 8000 Hz scanning speed (resonant scanner), frame averaging 4–6 times and z intervals of 0.5 µm. Z-stack projections (Sum-slices) were performed for each acquisition and signal and background were normalised using a custom script developed with the Fiji software (https://github.com/RDP-vbayle/SiCE_FIJI_Macro/blob/main/VariousSeedDev/MacroNormalisation-Creff_et_al_2024.ijm). For each set of data (folder), min and max pixel value are updated as following: Min pixel value (*i.e.* noise) is calculated in a 20 pixels square ROI in the upper left side of the SUM projections. Max pixel value corresponds to the maximum pixel value of all the SUM projection of the dataset (folder). To visualise seeds expressing tdTomato samples were excited with an LED emitting at 552 nm. The signal was collected at 580-650 nm.

For Feulgen staining, the LED excitation was set at 552 nm and the detector at 560-625 nm. Acquisition was performed with the 25x water immersion objective, 1.25x zoom, 20 % laser power, 8000 Hz scanning speed (resonant scanner), frame averaging 8 times and z intervals of 0.5 µm.

*In situ* hybridization

DNA templates for the probes used in *in situ* hybridizations were amplified using the primers listed in Table S1. Digoxigenin-labelled RNA probes were produced and hybridized to tissue sections following standard procedures. In brief, siliques were opened, fixed overnight in ice-cold PBS containing 4 % paraformaldehyde, dehydrated through an ethanol series, embedded in Paraplast Plus (Mc Cormick Scientific) and sectioned (8 µm). Immobilized sections were dewaxed and hydrated, treated with 2x saline sodium citrate (20 min), digested for 15 min at 37 °C with proteinase K (20 mg/mL) in 50 mM Tris-HCl, pH 7.5, 5 mM EDTA), treated for 2 min with 0.2

% glycine in PBS, rinsed, post-fixed with 4 % paraformaldehyde in PBS (10 min), rinsed, treated with 0.25 % w/v acetic anhydride in 100 mM triethanolamine (pH 8.0 with HCl) for 10 min, rinsed and dehydrated. Sections were then hybridized under coverslips overnight at 50 °C with RNA probes (produced using DIG RNA labelling kit (Roche)) diluted in DIG easy Hyb solution (Roche) following the manufacturer's instructions. Following hybridization, the slides were extensively washed in 0.1x saline sodium citrate and 0.5 % SDS at 50 °C (3 h), blocked for 1 h in 1 % blocking solution (Roche) in TBS and for 30 min in BSA solution (1 % BSA, 0.3 % Triton-X-100, 100 mM Tris-HCl, 100 mM NaCl, 50 mM $MgCl_2$), and then incubated in a 1/3000 dilution of in alkaline phosphatase-conjugated antidigoxigenin antibody (Roche) in BSA solution for 2 h at RT. Sections were extensively washed in BSA solution, rinsed and treated overnight in the dark with a buffered NBT/BCIP solution. Samples were rinsed in water before air drying and mounting in Entellan (Sigma). Imaging was carried out using a Zeiss Axioimager 2 equipped with a 20× dry objective under bright-field illumination.

Quantification of fertilization events from crosses with *kpl1-2* H2B:tdTomato

5 DAP siliques were opened and all the seeds were mounted in water between a slide and a coverslip. Venus/YFP and tdTomato signals of each seed were collected under UV illumination using a Zeiss Axioimager 2 equipped with a 20x dry objective.

Cytoplasmic calcium influx measurements

*Nicotiana benthamiana* plants stably expressing the Aequorin calcium reporter (*52*, *68*) were used to transiently express receptors of interest. In brief, constructs for the expression of receptors of interest were transformed into the *Agrobacterium tumefaciens* strain GV3101. Liquid L-media cultures containing 50 mg $mL^{-1}$ Kanamycin were inoculated with plate-grown colonies of *Agrobacterium* and grown at 28 °C overnight with shaking. The bacteria were pelleted by centrifugation and resuspended in 10 mM $MgCl_2$ to $O.D._{600} = 0.2$. This suspension was infiltrated into 4-week-old *N. benthamiana* plants. The following day, leaf disks were taken using 4-mm cutaneous biopsy punches with plungers (BPP-40F, KAI INDUSTRIES CO.,LTD) and floated on 100 µL of 20 µM coelenterazine (EC14031, Biosynth) in the dark overnight (*69*). The following morning, readings were taken using a VARIOSKAN MUTIPLATE READER (ThermoFisher) before and after the addition of 50 µL of 3x concentrated peptide solution or mock. Each leaf disk was discharged by the addition of 150 µL discharging solution (2 M $CaCl_2$ 20 % EtOH).

Western blotting

All steps were carried out on ice or at 4 °C and all buffers and tubes were precooled. Receptors of interest were transiently expressed in *Nicotiana benthamaia*, as described for calcium assays. Fourty-eight hours post-infiltration, ten ⌀ 4-mm leaf disks were flash -frozen in a 2mL tube with glass beads. Tissue was ground using a Genogrinder. Subsequently, 300 µL of protein extraction buffer (50 mM Tris pH 7.5, 150 mM NaCl, 2.5 mM EDTA, 10 % glycerol, 1 % IGEPAL, 5 mM DTT, 1 % plant protease inhibitor cocktail [P9599, Sigma]) was added to the tissue and incubated on a tube rotator. Samples were subsequently centrifuged at 13.2 *g* for 5 min before an aliquot of the supernatant was removed and denatured with 2x LDS Sample buffer (Merck). Samples were run on a polyacrylamide gel and transferred using the Trans-Blot Turbo Transfer system (Biorad). Western blotting was performed using α-GFP-HRP (sc-9996, Santa Cruz; 1:5000) and Coomassie brilliant blue (CBB), as a loading control.

Peptide synthesis

Peptide synthesis was performed by Genscript; specifics are outlined in Table S1.

Statistical analysis

Information regarding sample size and statistical tests can be found in each individual figure caption. The data were analysed using Excel (v. 2019), the R software (v.4.3.0), Graphpad Prism (v. 10.6.1 (799)) or Jasp (v.0.18.3). For box plot representation, the box represents the 25–75 % quartiles, and the median is represented by a horizontal line within the box. The whiskers of the box represent the minimal and maximal values of the sample. Contingency bar graph representation was used to represent quantification of phenotype repartition. Statistical analyses were conducted using Jasp (v.0.18.3 and v.0.19.3). For comparisons between two groups, a Student's *t*-test was used. For comparisons between multiple groups, a one-way ANOVA with Tukey's multiple comparison test was used. For RT-qPCR experiments, the groups were compared using a Kruskal-Wallis test followed by Dunn's Post-hoc test. Frequency comparisons were done with Chi-square tests.

Figure S1

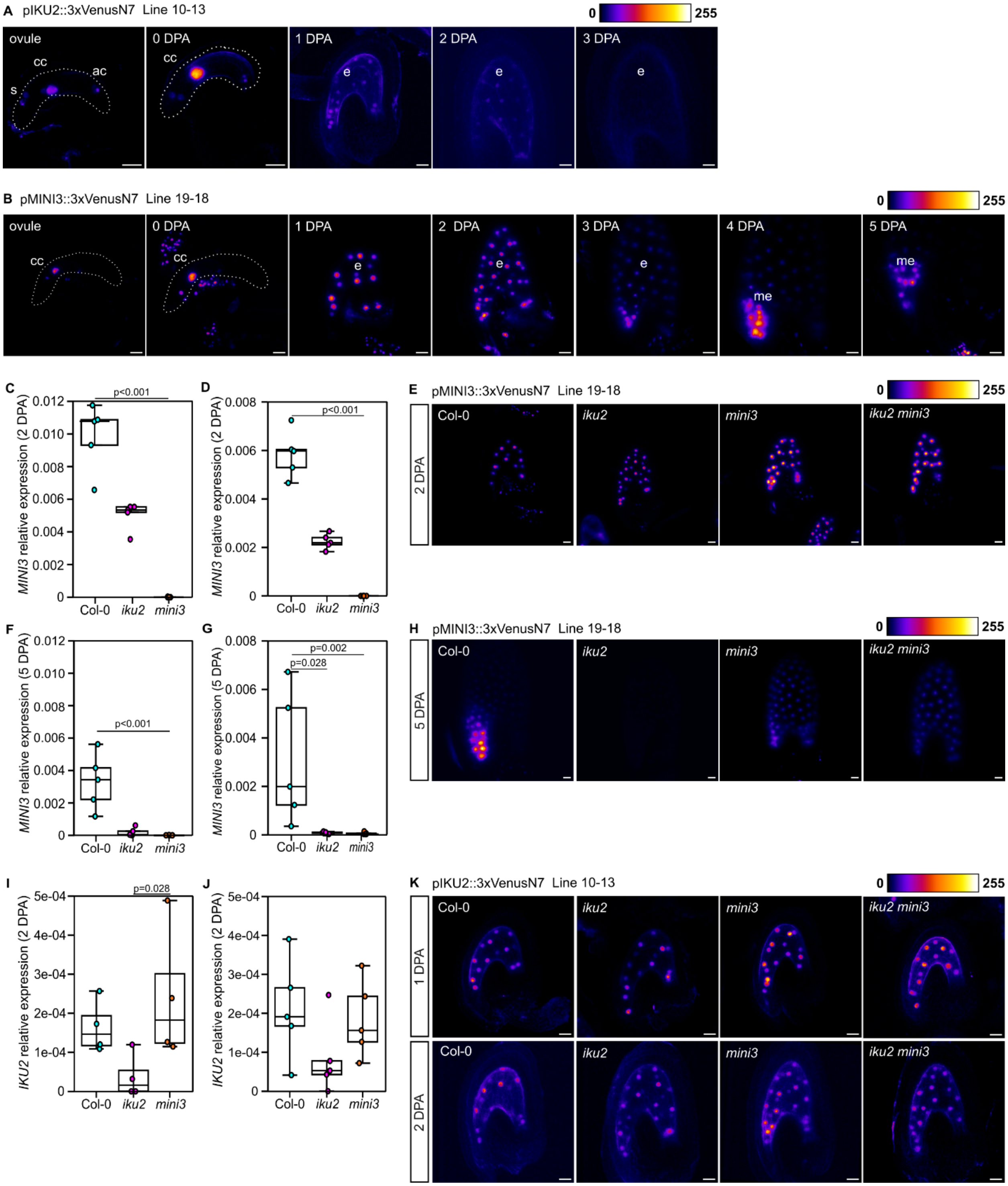


**Fig S1. IKU2 is required to maintain *MINI3* expression in the micropylar pole of the endosperm.** (Related to Fig 1)**.** (**A**) *IKU2* promoter activity in Col-0 background using a transcriptional reporter line during early seed development; s, synergid cells; cc, central cell; ac, antipodal cells; e, endosperm. Representative images from n = 7 seeds per time point. (**B**) *MINI3*

promoter activity in Col-0 background using transcriptional reporter line during early seed development; cc, central cell; e, endosperm; me, micropolar endosperm. Representative images from n = 6-8 seeds per time point. (**C,D**) Relative levels of *MINI3* transcript in 2 days post anthesis (DPA) siliques. (**E**) *MINI3* promoter activity in single and double mutant backgrounds at 2 DPA. Representative images from n = 10 seeds per genotype per time point. (**F,G**) Relative levels of *MINI3* transcript in 5 DPA siliques. (**H**) *MINI3* promoter activity in single and double mutant backgrounds at 5 DPA. n = 10 seeds per genotype per time point. (**I,J**) Relative levels of *IKU2* transcript in 2 DPA siliques. (**K**) *IKU2* promoter activity in single and double mutant background at 1 and 2 DPA. Representative images from n = 5-8 seeds per genotype per time point. Scale bars = 20 µm. (C-D-F-G-I-J) Two independent biological replicates of experiments shown in Fig 1 (D,E,G). In total this experiment was repeated 3 times. Transcript levels assessed by RT-qPCR. Each dot represents one plant (n = 5 samples in each case). p-values obtained using the Kruskal-Wallis followed with Dunn's Post-hoc test. (A,B,E,H,K) An independent biological replicate of data shown in Fig 1 (B,C,F,H)). In total this experiment was repeated 2 to 3 times in two independent transgenic lines and 11-34 seeds per genotype per time point were analysed.

Figure S2

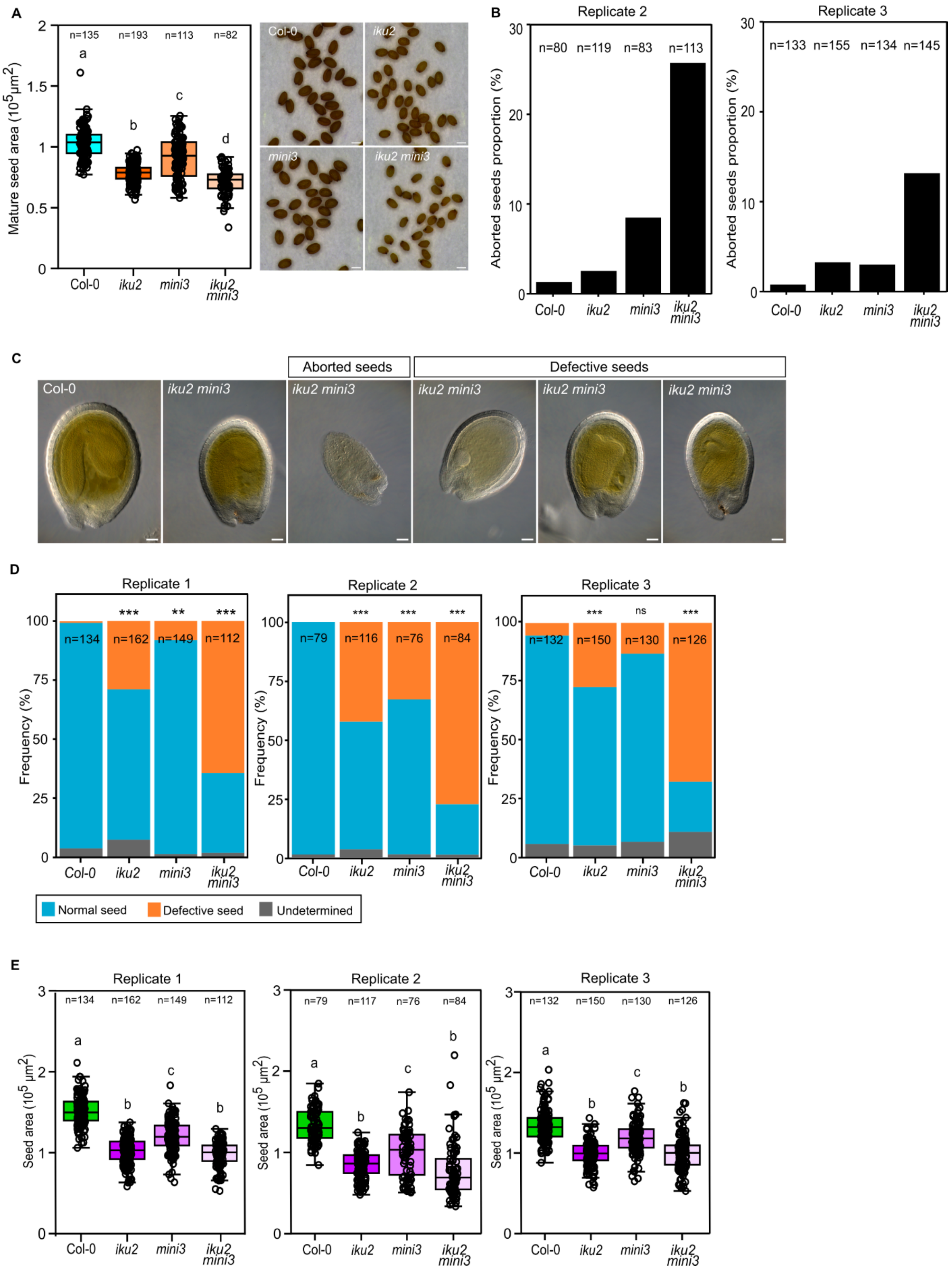
A
Mature seed area ($10^5 \mu m^2$)
n=135 n=193 n=113 n=82
a b c d
Col-0 iku2 mini3 iku2 mini3
B
Replicate 2
n=80 n=119 n=83 n=113
Replicate 3
n=133 n=155 n=134 n=145
Aborted seeds proportion (%)
C
Aborted seeds
Defective seeds
Col-0
iku2 mini3
D
Replicate 1
Replicate 2
Replicate 3
Frequency (%)
n=134 n=162 n=149 n=112
n=79 n=116 n=76 n=84
n=132 n=150 n=130 n=126
*** ** *** ns
Normal seed
Defective seed
Undetermined
E
Replicate 1
Replicate 2
Replicate 3
Seed area ($10^5 \mu m^2$)
n=134 n=162 n=149 n=112
n=79 n=117 n=76 n=84
n=132 n=150 n=130 n=126

**Fig S2. IKU2 and MINI3 act together to promote seed growth and normal seed development.** (Related to Fig 2). (**A**) Independent biological replicate of experiment shown in Fig 2A. Left: Measurements of Col-0, *iku2*, *mini3* and *iku2 mini3* mature seed areas from primary inflorescence stems. Total number of seeds analysed is indicated. Statistical groups established using ANOVA with Tukey's multiple comparison test; $p < 0.001$. Right: Representative images of dry seeds. Scale bars = 200 µm. (**B**) Quantification of aborted seeds in Col-0, *iku2*, *mini3* and *iku2 mini3*. The total number of seeds analysed is indicated. Two independent biological replicates of experiments shown in Fig 2 (C) are presented. In total this experiment was repeated 3 times and 347-443 seeds per genotype were analysed. (**C**) Representative Col-0 and *iku2 mini3* seeds at 8 DPA, illustrating the phenotypes of defective and aborted seeds quantified in (B) and (D,E). Scale bars = 50 µm. (**D**) Quantification of seed phenotypes in Col-0, *iku2*, *mini3* and *iku2 mini3* at 8 DPA. Three independent biological replicates are presented. Data are shown as contingency bar graphs, and the total number of seeds assessed per genotype is indicated. $X^2$ test; $^{***}p < 0.00$, ns: not significant. (**E**) Measurements of Col-0, *iku2*, *mini3* and *iku2 mini3* developing seed areas at 8 DPA from primary inflorescence stems. Three independent biological replicates are presented. Total number of seeds analysed is indicated. Statistical groups established using ANOVA with Tukey's multiple comparison test; $p < 0.001$.

### Figure S3

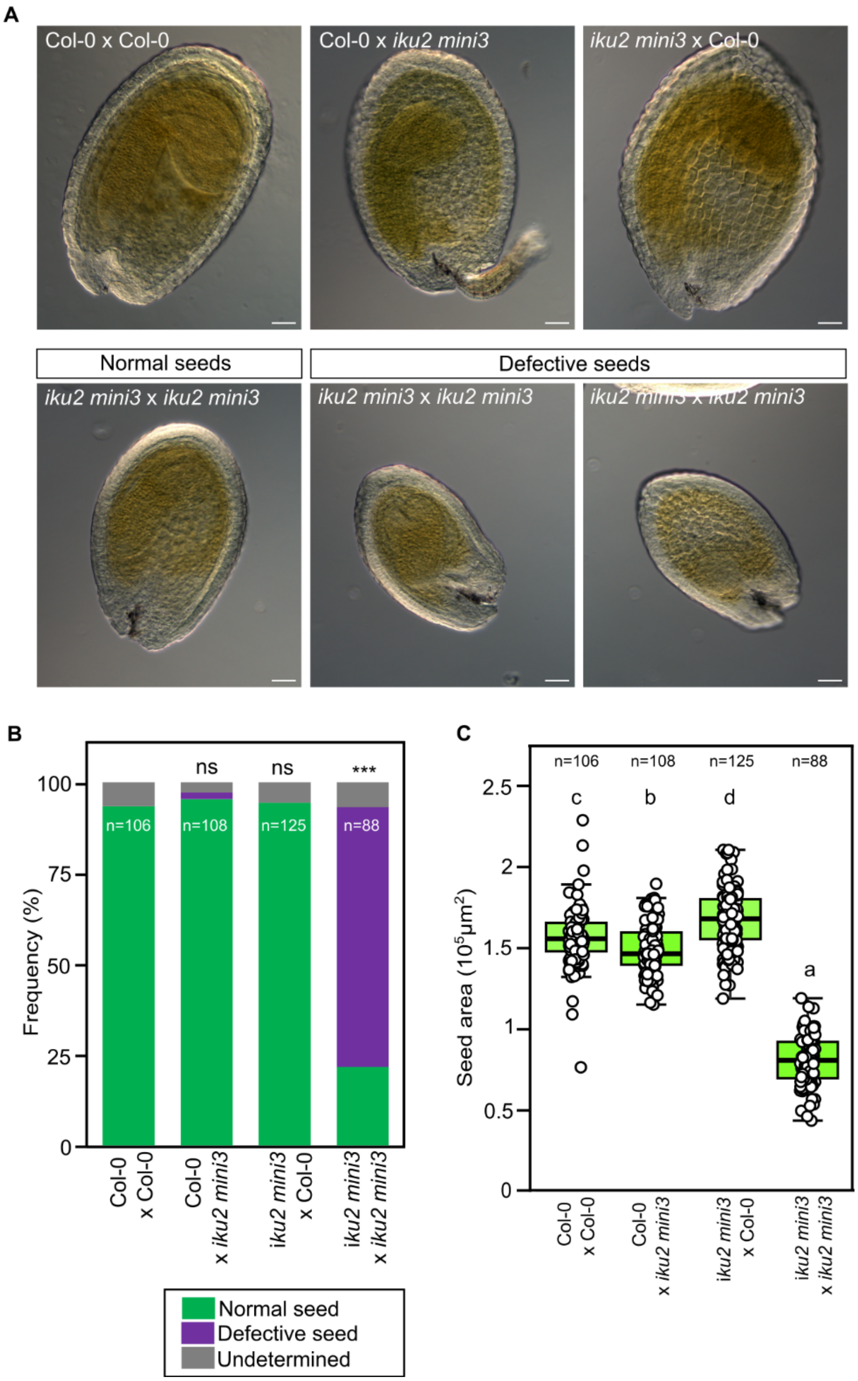


**Fig S3. The *iku2 mini3* phenotype is dependent on the zygotic genotype.** (Related to Fig 2). (**A**) Representative seeds from F1 siliques at 8 DPA, illustrating the phenotypes quantified in (B) and (C). Scale bars = 50 µm. (**B**) Quantification of seed phenotypes from F1 siliques at 8 DPA. Data are shown as a contingency bar graphs, and the total number of seeds assessed per genotype is indicated. $X^2$ test; ***$p < 0.001$, ns: not significant. (**C**) Measurement of seed areas from F1 siliques. Statistical groups established using ANOVA with Tukey's multiple comparison test; $p < 0.001$ between all groups except a and b ($p=0.012$). (B,C) Seeds from three independent crosses were pooled.

Figure S4

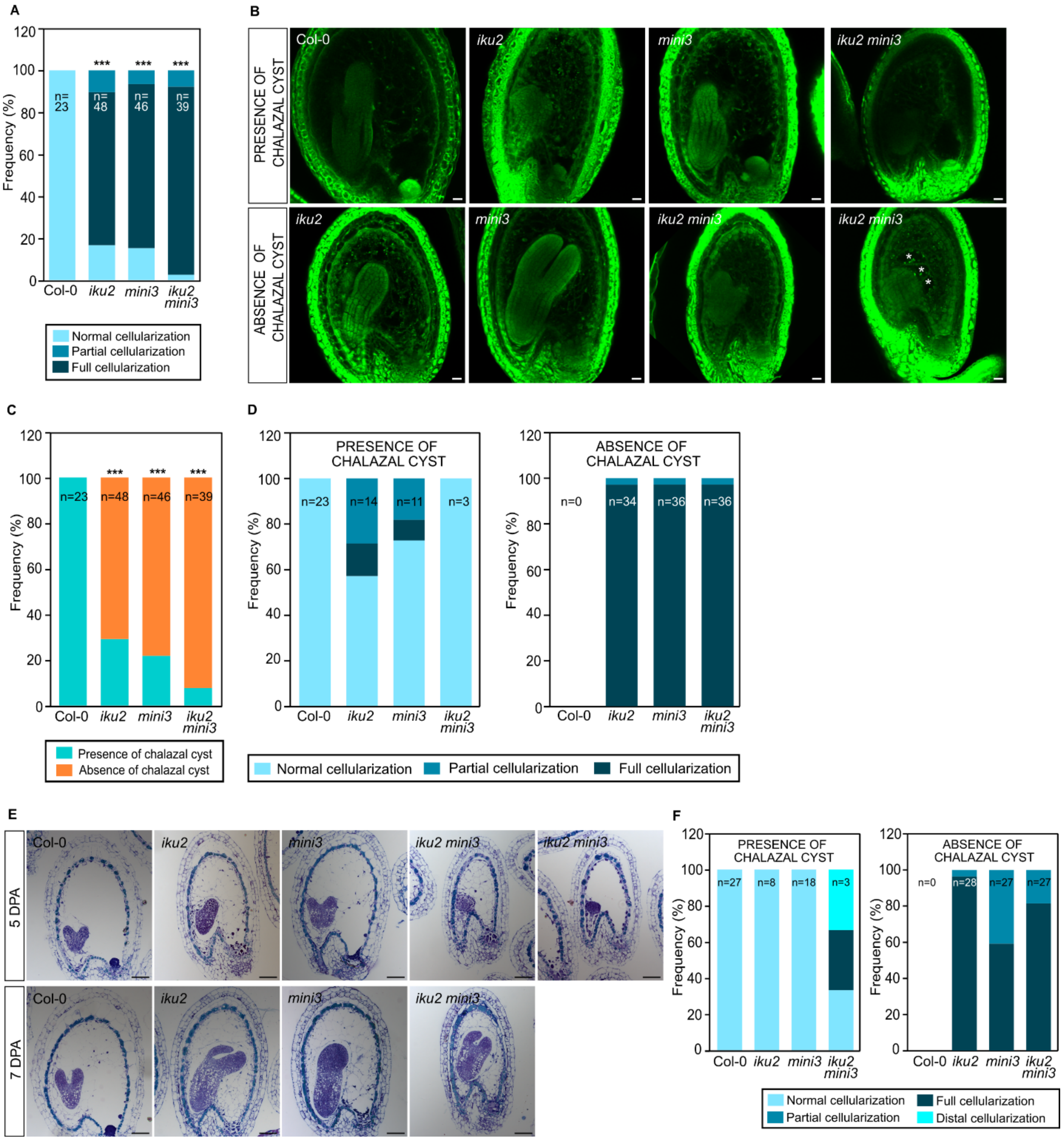


**Fig S4. Endosperm development is severely compromised in HAIKU pathway mutants.** (Related to Fig 2). (**A**) Patterns of endosperm cellularization in Col-0, *iku2*, *mini3* and *iku2 mini3* 7 DPA seeds assessed with Feulgen staining. Data are shown as a contingency bar graph, and the total number of seeds per genotype is indicated. $X^2$ test. ***p < 0.001. (**B**) Representative images of Col-0 and mutant seeds stained with Feulgen protocol at 7 DPA. White stars indicate uncellularized endosperm in an unexpected position. Scale bars = 20 µm. (**C**) Quantification of chalazal cyst presence in Col-0, *iku2, mini3* and *iku2 mini3* seeds obtained from Feulgen staining.

Data are shown as a contingency bar graph, and the total number of seeds per genotype is indicated. $X^2$ test. ***p < 0.001. (**D**) Endosperm cellularization as a function of chalazal cyst presence in Col-0, *iku2, mini3* and *iku2 mini3* seed. Data are shown as contingency bar graphs, and the total number of seeds analysed per genotype is indicated. (**E**) Representative images from an independent biological replicate of the experiment shown in Fig 2B. In total this experiment was repeated 2 times and 11 to 18 seeds per genotype per time point were analysed. Median sections of Col-0 and mutant seeds stained with toluidine blue at two stages of development. Scale bars = 50 µm. (**F**) Endosperm cellularization as a function of chalazal cyst presence in Col-0, *iku2, mini3* and *iku2 mini3* seeds. Data are shown as a contingency bar graph, and the total number of seeds analysed per genotype is indicated.

Figure S5

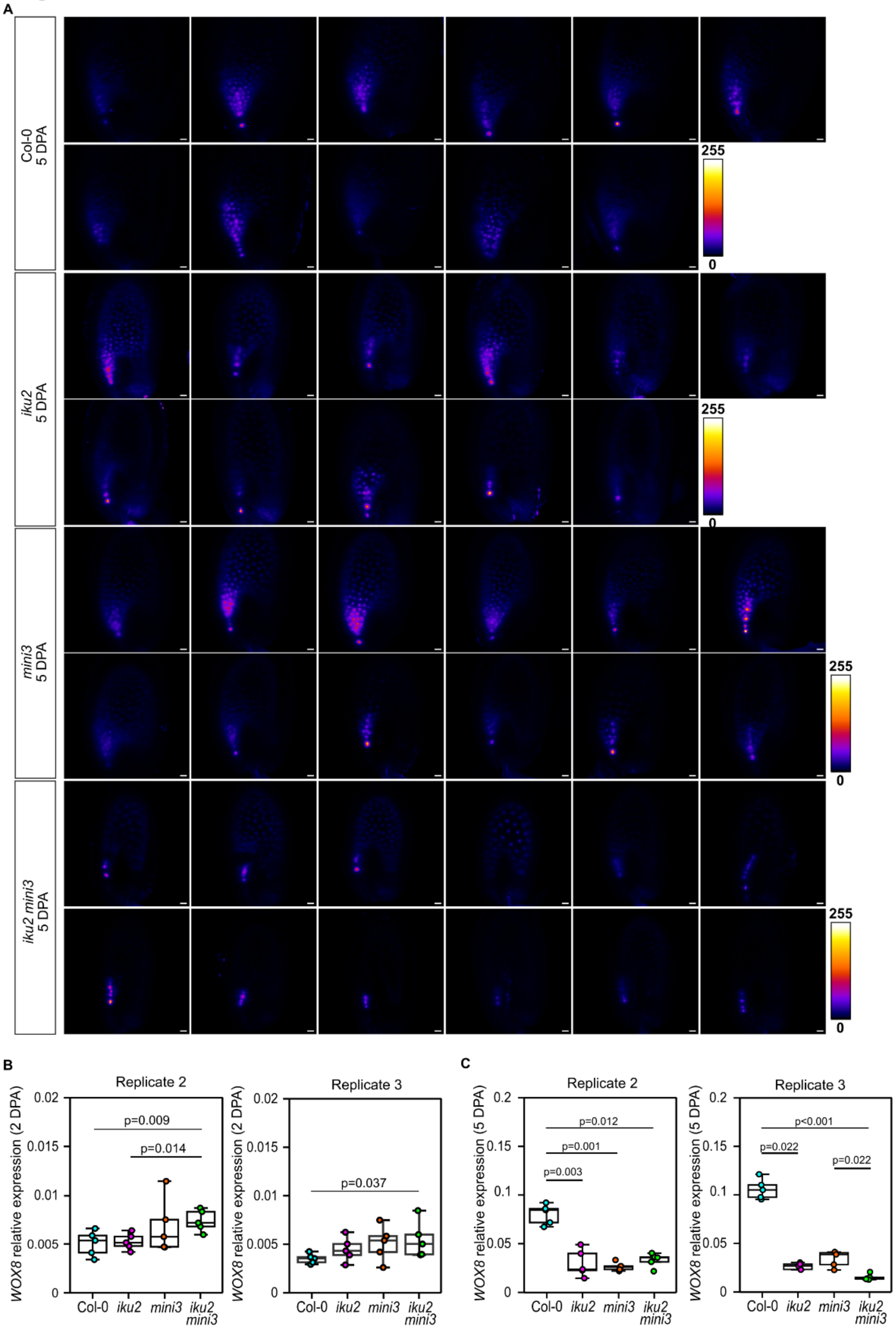


**Fig S5. The HAIKU pathway regulates the expression of *WOX8* in the micropylar endosperm.** (Related to Fig 2). (**A**) Global view of *WOX8* promoter activity in Col-0, *iku2*, *mini3* and *iku2 mini3* at 5 DPA. All images from one experiment are shown. In total this experiment was repeated 2 times and 21 to 23 seeds were analysed. Scale bars = 20 µm. (**B**) Relative levels of *WOX8* transcript in 2 DPA siliques. (**C**) Relative levels of *WOX8* transcript in 5 DPA siliques. (B,C) Two independent biological replicates of experiments shown in Fig 2 (H,I) are presented. In total this experiment was repeated 3 times. Transcript levels assessed by RT-qPCR. Each dot represents one plant (n = 5 samples in each case). p-values obtained using the Kruskal-Wallis test followed by Dunn's Post-hoc test.

Figure S6

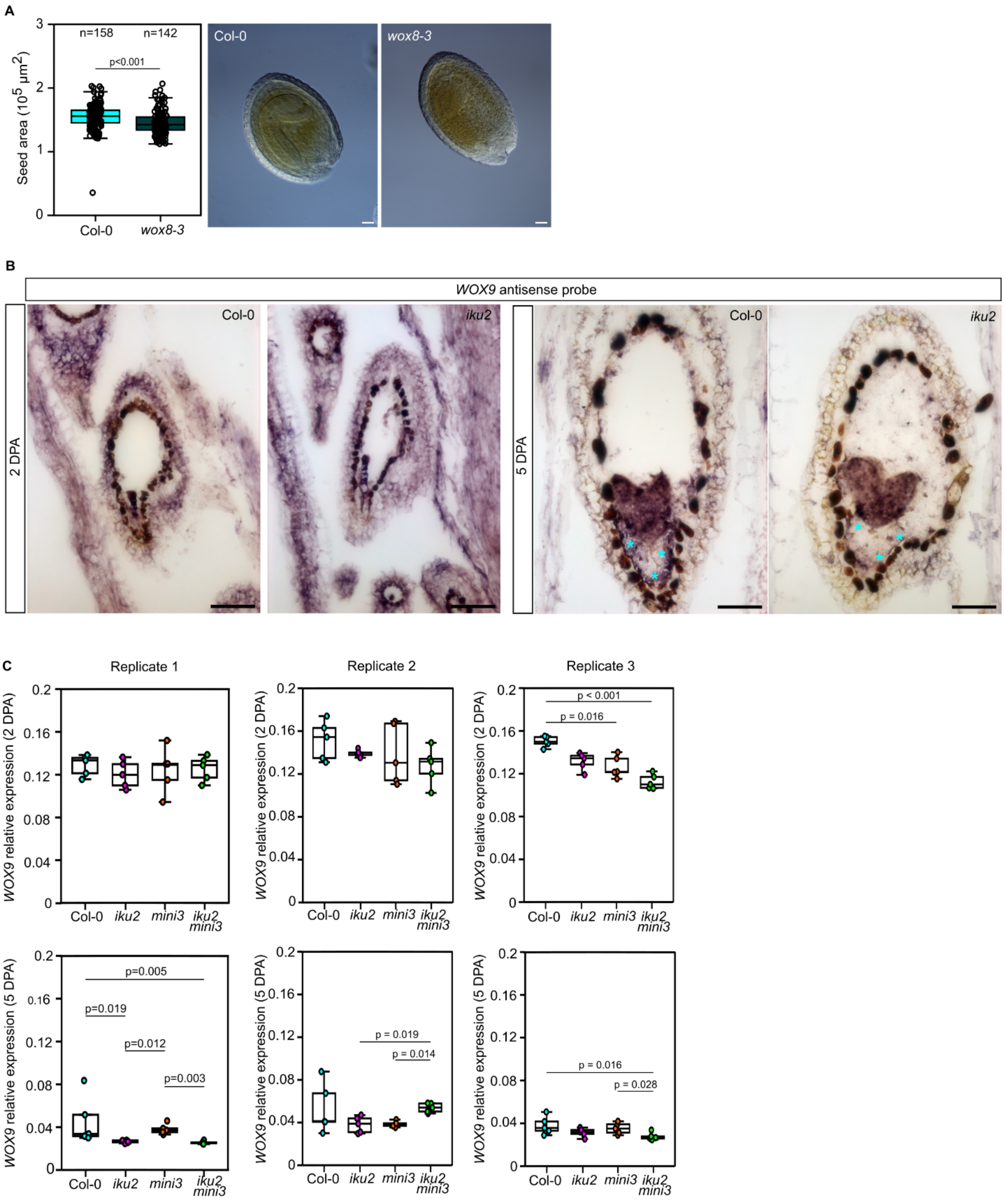


**Fig S6. The HAIKU pathway acts to establish endosperm polarity through the activation of *WOX8* and *WOX9* in the endosperm.** (Related to Fig 2). (**A**) Left: Measurement of the areas of developing seeds from Col-0 and *wox8-3* primary inflorescence stems at 8 DPA. Total number of seeds is indicated. p-values were established using Student's t-test. Right: Representative images of Col-0 and *wox8-3* seeds at 8 DPA. Scale bars = 50 µm. (**B**) Representative images of 2 and 5 DPA paraplast embedded seed sections hybridized with *WOX9* antisense probe. Micropylar endosperm signal is indicated with cyan stars. (**C**) Relative levels of *WOX9* transcript in 2 and 5 DPA siliques assessed by RT-qPCR in three independent experiments. Each dot represents one plant (n = 5 samples in each case). Statistical differences established using the Kruskal-Wallis test followed by Dunn's Post-hoc test.

Figure S7

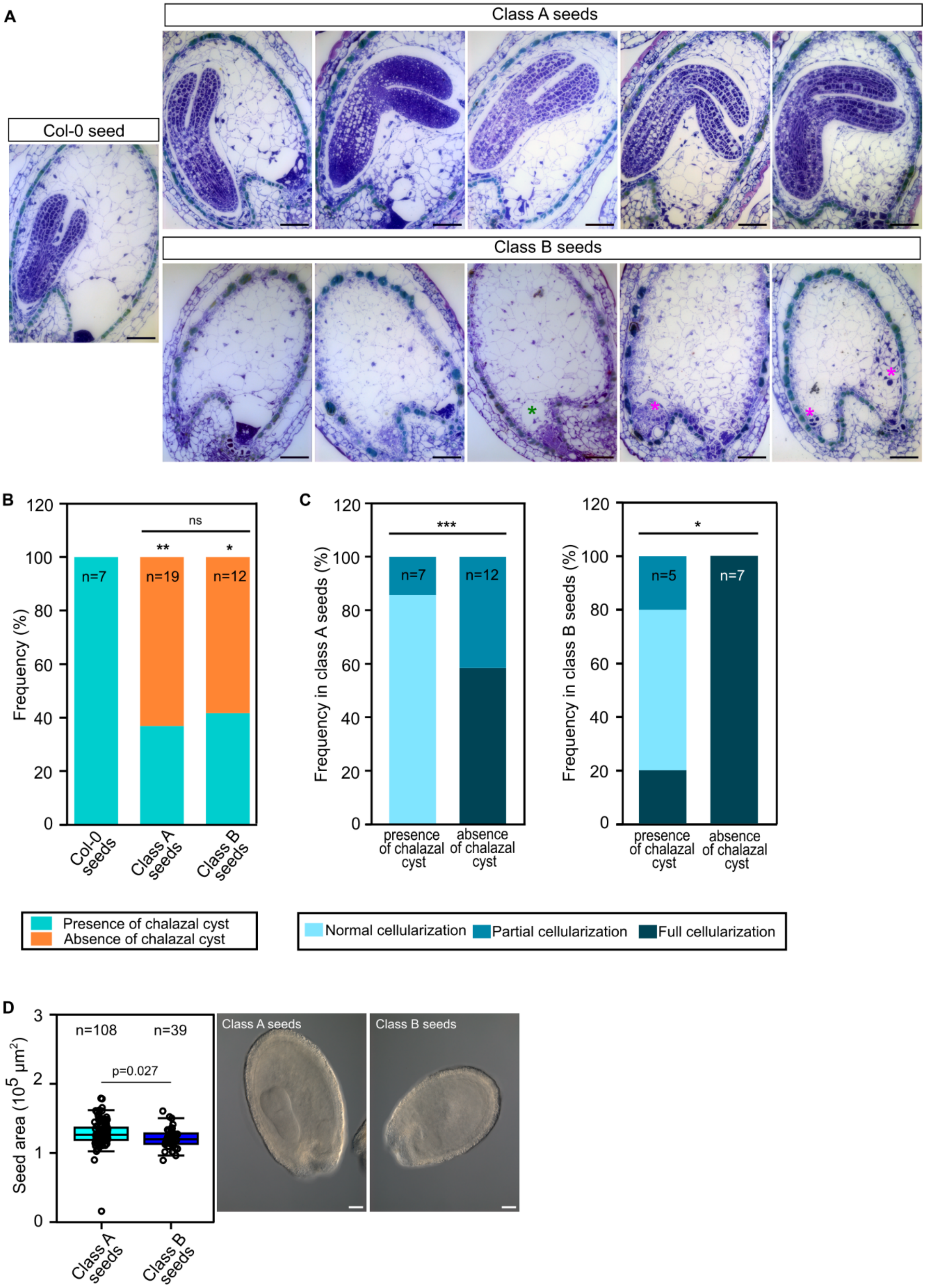
A
Class A seeds
Col-0 seed
Class B seeds
B
Frequency (%)
120
100
80
60
40
20
0
ns
** *
n=7 n=19 n=12
Col-0 seeds
Class A seeds
Class B seeds
Presence of chalazal cyst
Absence of chalazal cyst
C
Frequency in class A seeds (%)
***
n=7 n=12
presence of chalazal cyst
absence of chalazal cyst
Frequency in class B seeds (%)
*
n=5 n=7
presence of chalazal cyst
absence of chalazal cyst
Normal cellularization
Partial cellularization
Full cellularization
D
Seed area (10^5 µm^2)
3
2
1
0
n=108 n=39
p=0.027
Class A seeds
Class B seeds
Class A seeds
Class B seeds

**Fig S7. The HAIKU pathway acts to establish endosperm polarity through the activation of *WOX8* and *WOX9* in the endosperm.** (Related to Fig 2). (**A**) Representative images of Col-0 and *wox8 wox9*/+ seeds stained with toluidine blue at 7 DPA. Potential over-proliferation of nucellus-like tissues is indicated by magenta stars. The green star shows the lack of cellularization above the mutant embryo. (**B**) Quantification of chalazal cyst presence in Col-0 and class A and class B (containing *wox8 wox9* double mutant embryos and endosperms) seeds from self-pollinated *wox8 wox9*/+ plants obtained by toluidine blue staining of sections at 7 DPA. Data are shown as a contingency bar graph, and the total number of seeds analysed per class is indicated. Please note that class B seeds represent about 25% of seeds produced. $X^2$ test; **$p < 0.01$, *$p < 0.05$, ns: not significant. (**C**) Endosperm cellularization pattern as a function of the presence of the chalazal cyst in class A and class B seeds obtained with toluidine blue staining. Data are shown as a contingency bar graphs, and the total number of seeds analysed per class is indicated. $X^2$ test; ***$p < 0.001$, *$p < 0.05$. (**D**) Left: Measurement of class A and class B seed areas from primary inflorescence stems at 6 DPA. Total number of seeds is indicated. p-values established using Student's t-test. Right: Representative images of class A and class B seeds at 6 DPA. Scale bars = 50 µm.

#### Figure S8

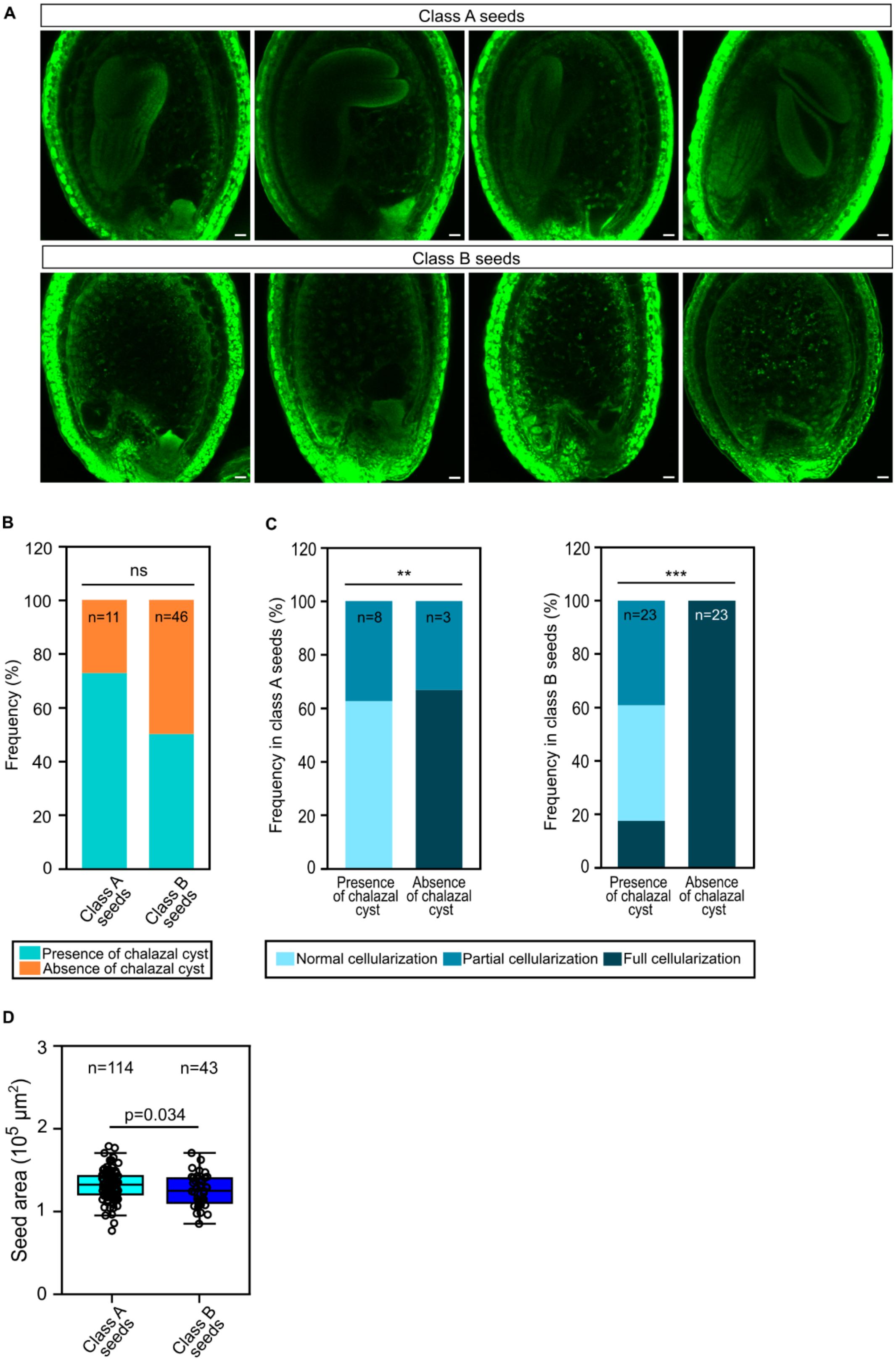
A
Class A seeds
Class B seeds
B
ns
n=11
n=46
Frequency (%)
Class A seeds
Class B seeds
Presence of chalazal cyst
Absence of chalazal cyst
C
**
n=8
n=3
Frequency in class A seeds (%)
***
n=23
n=23
Frequency in class B seeds (%)
Presence of chalazal cyst
Absence of chalazal cyst
Normal cellularization
Partial cellularization
Full cellularization
D
n=114
n=43
p=0.034
Seed area ($10^5$ μm$^2$)
Class A seeds
Class B seeds

**Fig S8. WOX8 and WOX9 act together to ensure normal endosperm development.** (Related to Fig 2). (**A**) Representative images of *wox8 wox9/+* 7 DPA seeds obtained with Feulgen staining. Scale bars = 20 µm. (**B**) Quantification of chalazal cyst presence in class A and class B (containing *wox8 wox9* double mutant embryos and endosperms) seeds from self-pollinated *wox8 wox9/+* plants obtained by Feulgen staining at 7 DPA. Data are shown as a contingency bar graph, and the total number of seeds analysed per class is indicated. Please note that class B seeds represent about 25 % of seeds produced by these crosses. $X^2$ test; ns: not significant. (**C**) Endosperm cellularization pattern as a function of the presence of the chalazal cyst in class A and class B seeds obtained with Feulgen staining. Data are shown as contingency bar graphs, and the total number of seeds analysed per class is indicated. $X^2$ test; $^{**}p < 0.05$, $^{***}p < 0.001$. (**D**) Independent biological replicate of the experiment shown in Figure S7. Measurements of class A and class B seed areas from primary inflorescence stems at 6 DPA. Total number of seeds is indicated. p-values established using Student's t-test.

### Figure S9

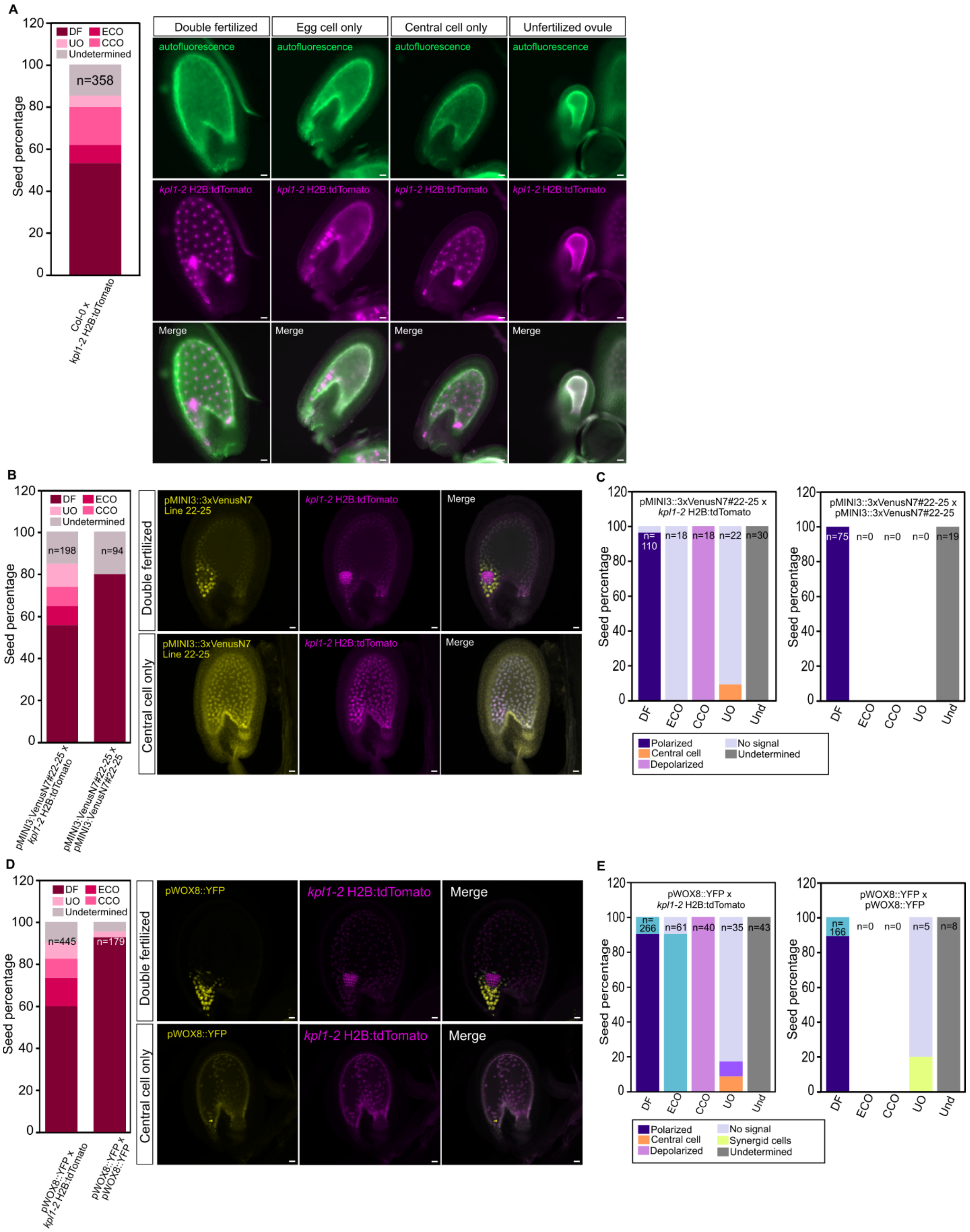
A
B
C
D
E
Seed percentage
DF ECO UO CCO Undetermined
n=358
Col-0 x kpl1-2 H2B:tdTomato
Double fertilized
Egg cell only
Central cell only
Unfertilized ovule
autofluorescence
kpl1-2 H2B:tdTomato
Merge
n=198
n=94
pMINI3::VenusN7#22-25 x kpl1-2 H2B:tdTomato
pMINI3::VenusN7#22-25 x pMINI3::VenusN7#22-25
pMINI3::3xVenusN7 Line 22-25
Polarized
Central cell
Depolarized
No signal
Undetermined
Synergid cells
pMINI3::3xVenusN7#22-25 x kpl1-2 H2B:tdTomato
pMINI3::3xVenusN7#22-25 x pMINI3::3xVenusN7#22-25
n=445
n=179
pWOX8::YFP x kpl1-2 H2B:tdTomato
pWOX8::YFP x pWOX8::YFP
pWOX8::YFP
DF ECO CCO UO Und

**Fig S9. A signal produced by the fertilized zygote or very young embryo is necessary for endosperm polarity establishment: Impact on *MINI3* and *WOX8* expression.** (Related to Fig 3). (**A**) Left: Quantification of fertilization events from crosses of Col-0 with *kpl1-2* H2B:tdTomato at 5 DAP. Right: Representative images of each category. (**B**) Left: Quantification of fertilization events from crosses of a second independent pMINI3::3xVenusN7 line (line 22-25) with *kpl1-2* H2B:tdTomato at 5 DAP. Total number of seeds is indicated. Right: Representative images of *MINI3* promoter activity in double fertilized seeds and cco seeds after Clearsee Alpha treatment at 5 DPA (line 22-25). (**C**) Left: Quantification of signal localization in line #22-25 depending upon fertilization events. Right: Quantification of signal localization depending upon fertilization events in self-pollination of *MINI3* reporter line (line 22-25). Total number of seeds is indicated. (B,C) Replicate of the experiment shown in Fig 3A using an independent transgenic line. (**D**) Left: Quantification of signal localization depending upon fertilization events. Total number of seeds analysed is indicated. Right: Representative images of *WOX8* promoter activity in double fertilized seeds and cco seeds after Clearsee Alpha treatment at 5 DAP. **(E**) Left: Quantification of fertilization events from crosses of pWOX8::YFP with *kpl1-2* H2B:tdTomato at 5 DAP. Total number of seeds analysed is indicated. Right: Quantification of signal localization depending upon fertilization events in self- pollination of *WOX8* reporter line. Total number of seeds analysed is indicated. (A,E) DF, double fertilized; ECO, egg cell fertilized only ; UO, unfertilized ovules; CCO, central cell fertilized only. Undetermined category corresponds to damaged seeds or seeds with signals that are difficult to distinguish. Scale bars = 20 μm.

#### Figure S10

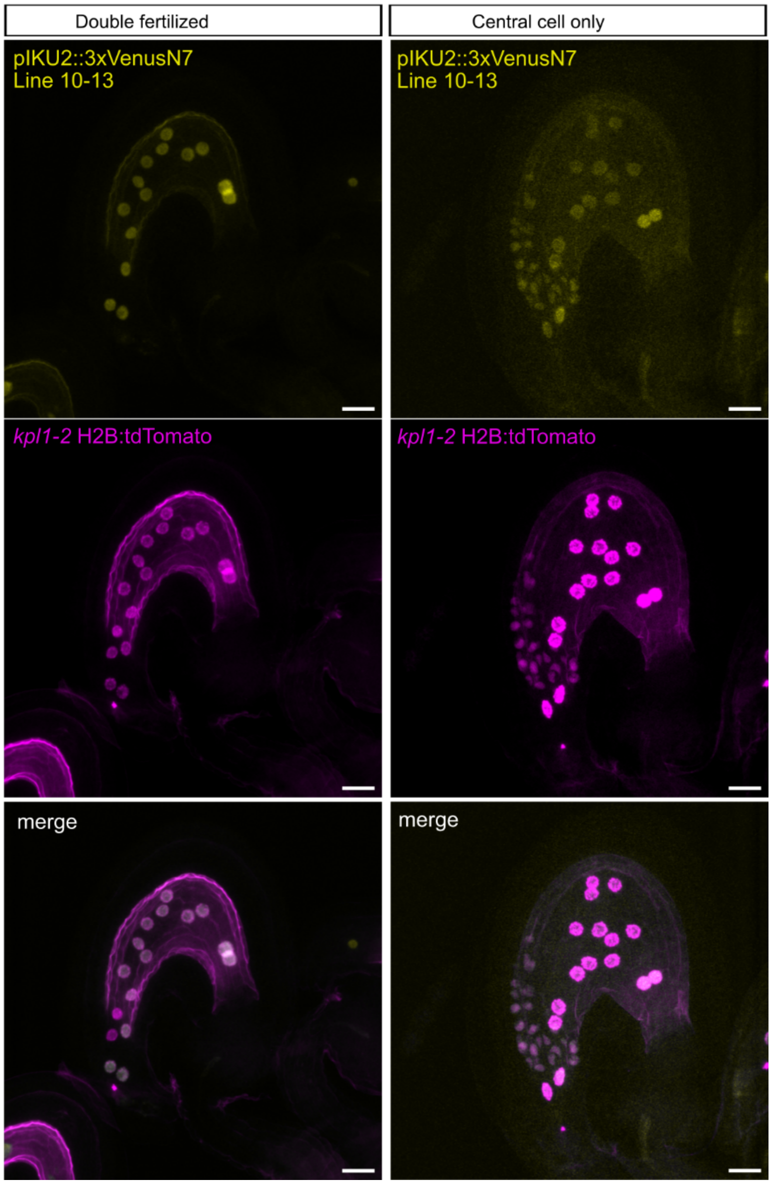


**Fig S10. A signal produced by the fertilized zygote or very young embryo is necessary for endosperm polarity establishment: Impact on *IKU2* expression.** (Related to Fig 3). Representative images of *IKU2* promoter activity in double and CCO fertilization events after Clearsee Alpha treatment at 2 DAP. Scale bars = 20 µm.

### Figure S11

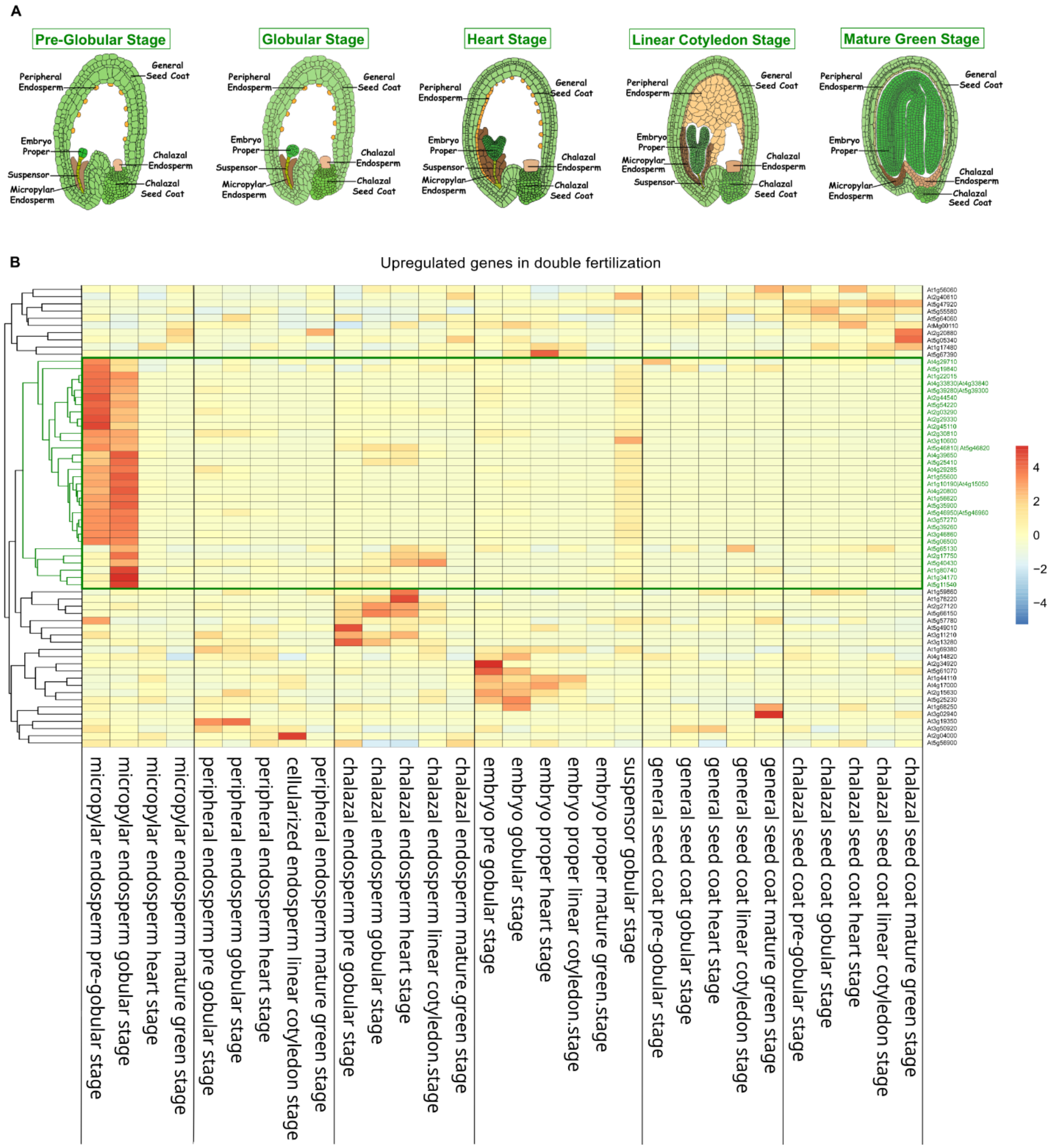


**Fig S11. A signal produced by the fertilized zygote or very young embryo is necessary for endosperm polarity establishment: Transcriptomic impact of loss of IKU2 function**. (Related to Fig 3). (**A**) Arabidopsis seed tissues at different stages of development. (http://seedgenenetwork.net/arabidopsis). (**B**) Heatmap representation of genes showing reduced expression in *iku2* seeds compared to wild-type seeds at 2DPA.

### Figure S12

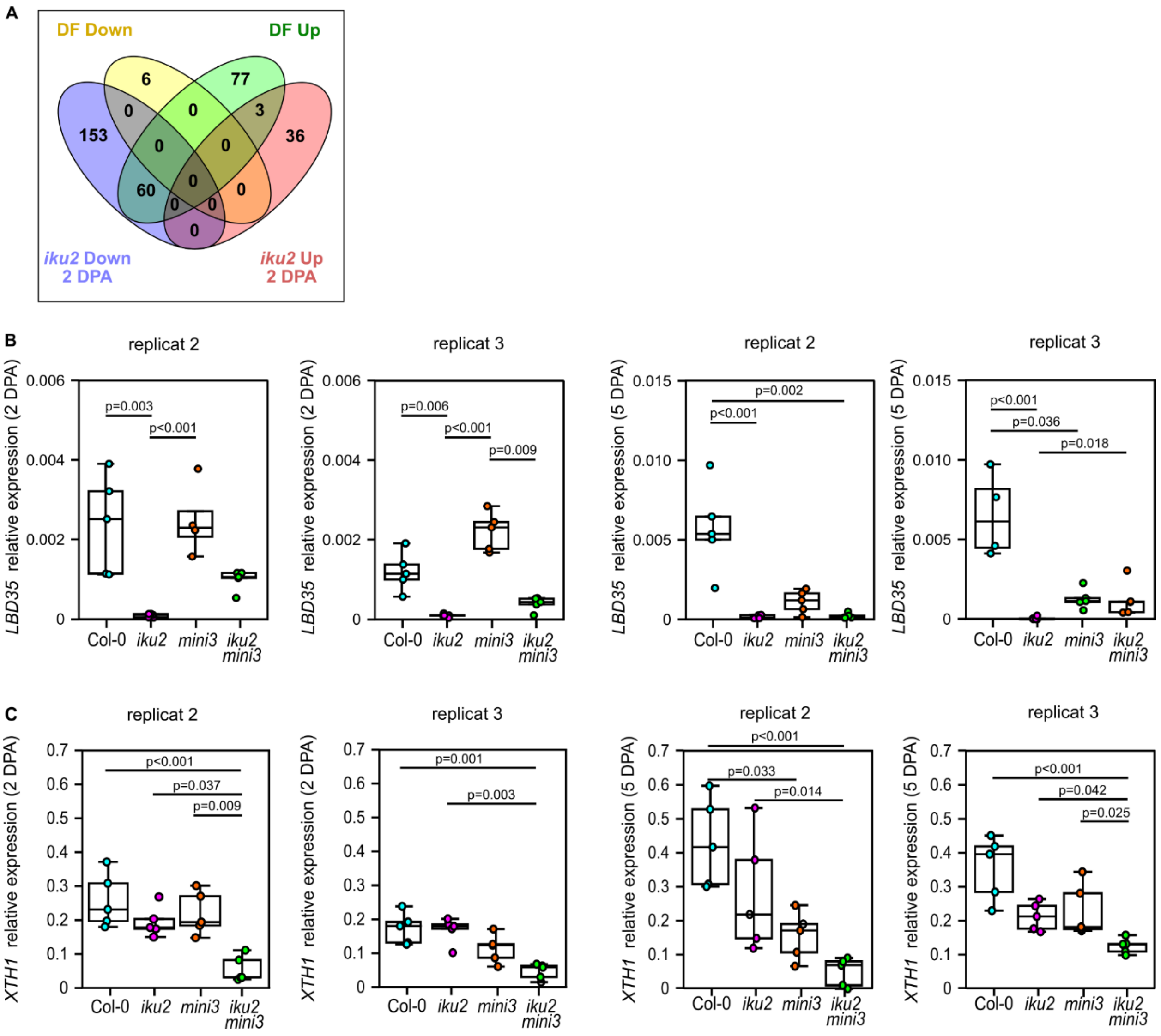


**Fig S12. A signal produced by the fertilized zygote or very young embryo is necessary for endosperm polarity establishment: Transcriptomic impact**. (Related to Fig 3). (**A**) Overlap between genes misregulated in *iku2* mutants at 2 DPA and genes misregulated specifically in double-fertilized seeds at 4 DPA. (**B**) Relative levels of *LBD35* transcript in 2 and 5 Days Post Anthesis (DPA) siliques. (**C**) Relative levels of *XTH1* transcript in 2 and 5 DPA siliques. (B,C) Two independent biological replicates of experiments shown in Fig 3 (D,E,F,G). In total this experiment was repeated 3 times. Transcript levels assessed by RT-qPCR. Each dot represents one plant (n = 5 samples in each case). p-values obtained using the Kruskal-Wallis test followed with Dunn's Post-hoc test.

Figure S13

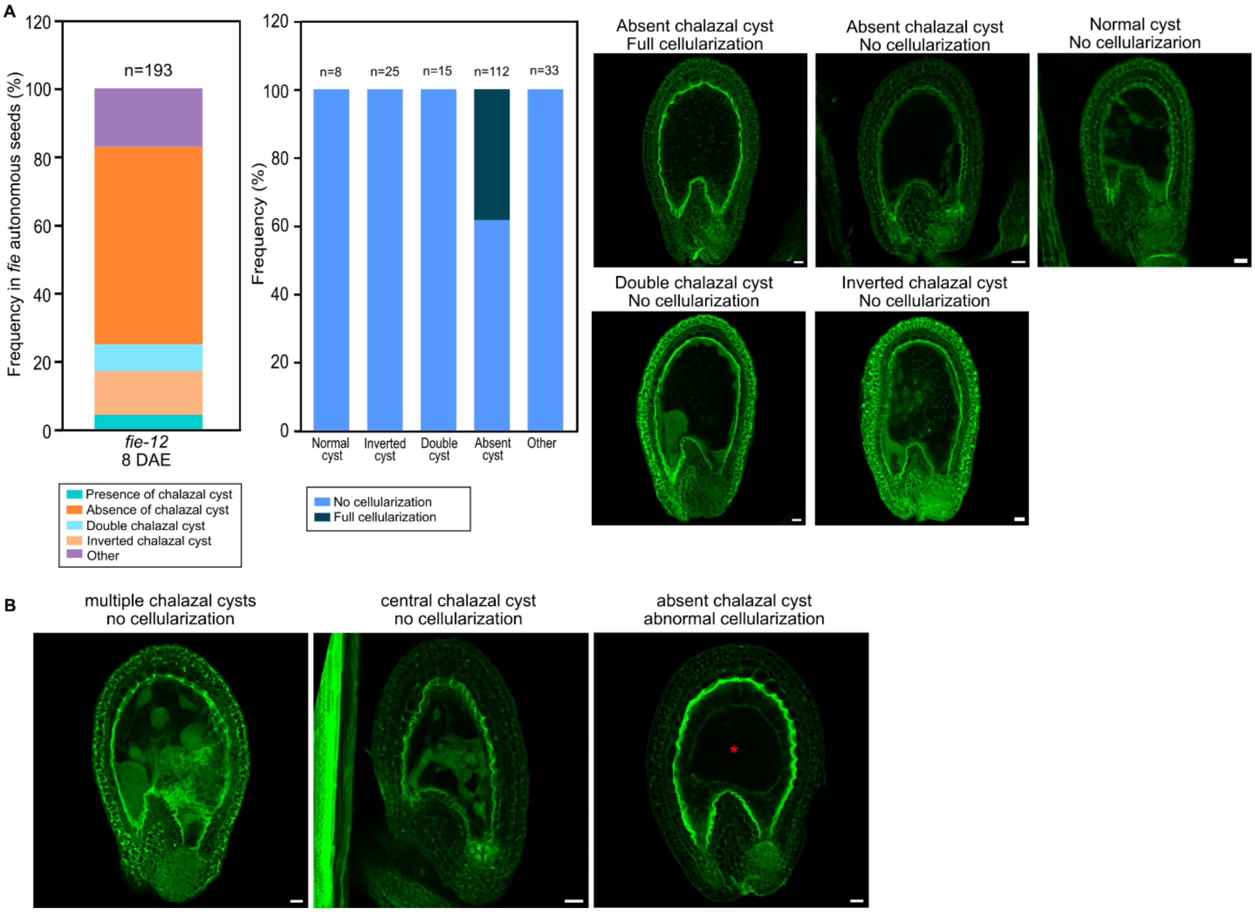


**Fig S13. A signal produced by the fertilized zygote or very young embryo is necessary for endosperm polarity establishment: Data relating to *fie* mutant autonomous endosperm development.** (Related to Fig 3). (**A**) Left: Presence and position of chalazal cyst in *fie* autonomous seeds at 8 DAE. Total number of seeds analysed is indicated. Right: Representative images of *fie* autonomous seeds harbouring various endosperm phenotypes at 8 DAE. (**B**) Representative images of 8 DAE *fie* autonomous seeds showing other types of cyst phenotype rarely observed; presence of several cysts in the endosperm cavity; presence of a big central cyst; abnormal cellularization in absence of chalazal cyst. Red star indicates an uncellularized zone in the center of the endosperm. (A,B) Scale bar = 20 µm.

### Figure S14

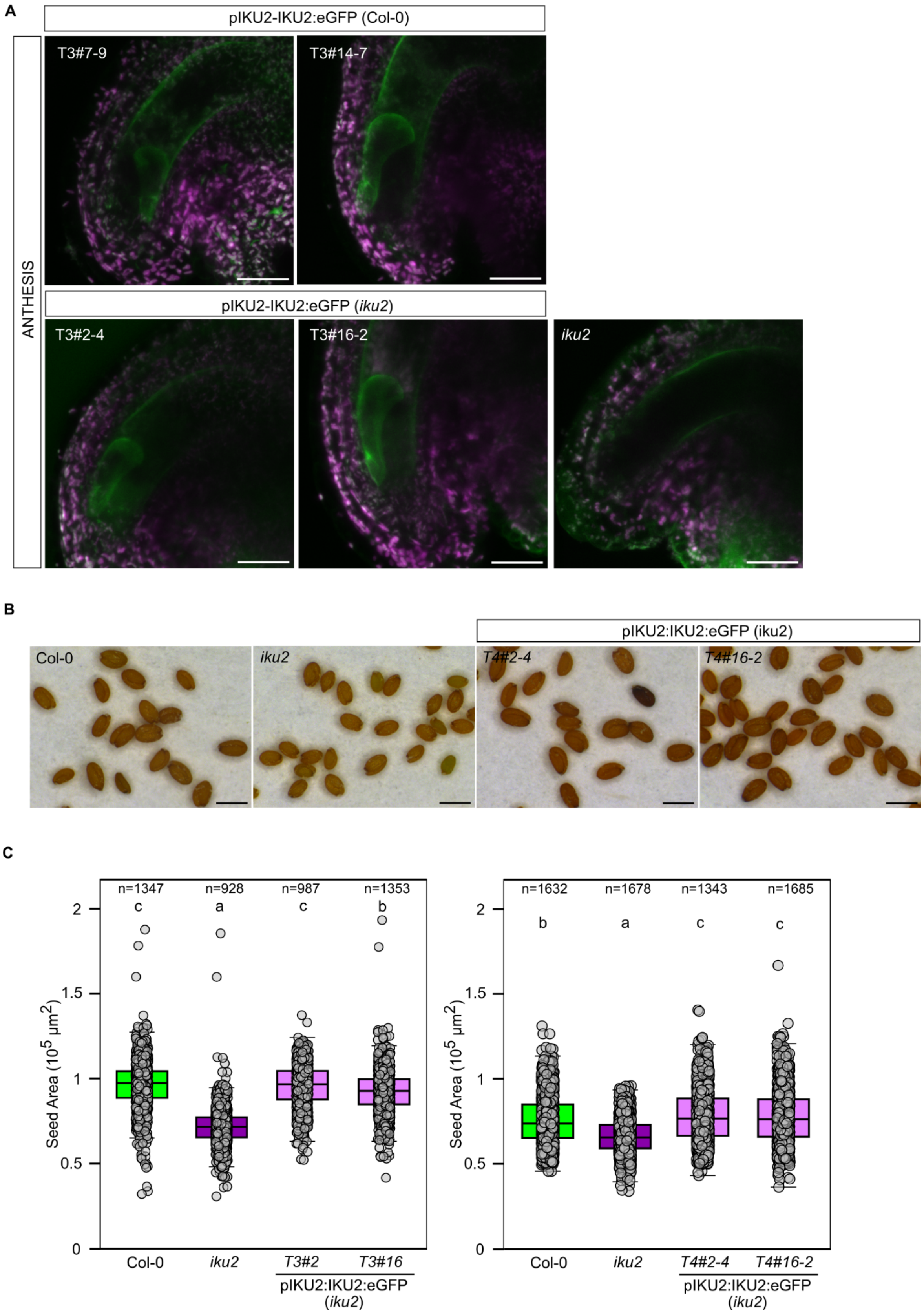
A
pIKU2-IKU2:eGFP (Col-0)
ANTHESIS
T3#7-9
T3#14-7
pIKU2-IKU2:eGFP (iku2)
T3#2-4
T3#16-2
iku2
B
pIKU2:IKU2:eGFP (iku2)
Col-0
iku2
T4#2-4
T4#16-2
C
n=1347
n=928
n=987
n=1353
c
a
c
b
Seed Area (10^5 μm^2)
2
1.5
1
0.5
0
Col-0
iku2
T3#2
T3#16
pIKU2:IKU2:eGFP
(iku2)
n=1632
n=1678
n=1343
n=1685
b
a
c
c
T4#2-4
T4#16-2

**Fig S14. A translational fusion of IKU2 to eGFP complements the iku2 phenotype.** (Related to Fig 3). (**A**) IKU2 localisation in plants expressing a complementing *pIKU2-IKU2:eGFP* construct in wild type and *iku2* plants. Representative images at anthesis stage. Scale bars = 20 µm. (**B**) Representative images of dry seeds from complemented *iku2* line. Scale bars = 200 µm. (**C**) Measurements of Col-0, *iku2* and complemented lines mature seed areas from primary inflorescence stems. (B,C) Seed sizes of two independent complemented lines. Experiments correspond to T3 and T4 generations respectively. Total number of seeds analysed is indicated. Statistical groups established using ANOVA with Tukey's multiple comparison test; $p < 0.001$.

Figure S15

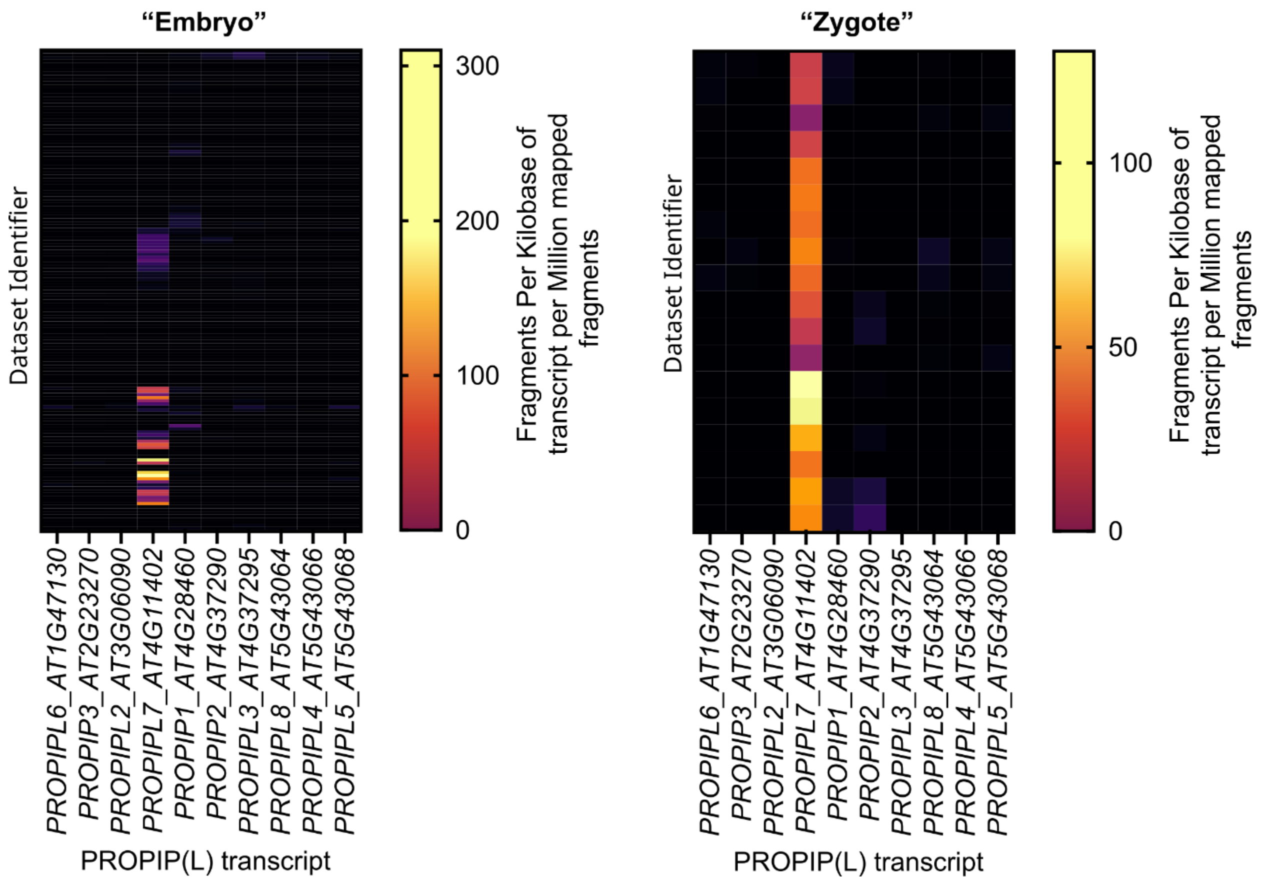


**Fig S15. Expression analysis of *PROPIP* and *PROPIPL* genes.** (Related to Fig 4). RNA expression data was extracted for annotated PIP(L)s from https://plantrnadb.com/athrdb/. Datasets with tissue metadata annotations of "embryo" and "zygote" were extracted and sorted according to dataset identifier (no biological relevance) and displayed as a heatmap. *PROPIPL1* was excluded from this analysis as alignment of the active epitopes suggests PIPL1 is more related to CEP peptides (CEP-like) (Fig S16), in accordance with its annotation on UNIPROT as CEP16 (Q5S502).

Figure S16

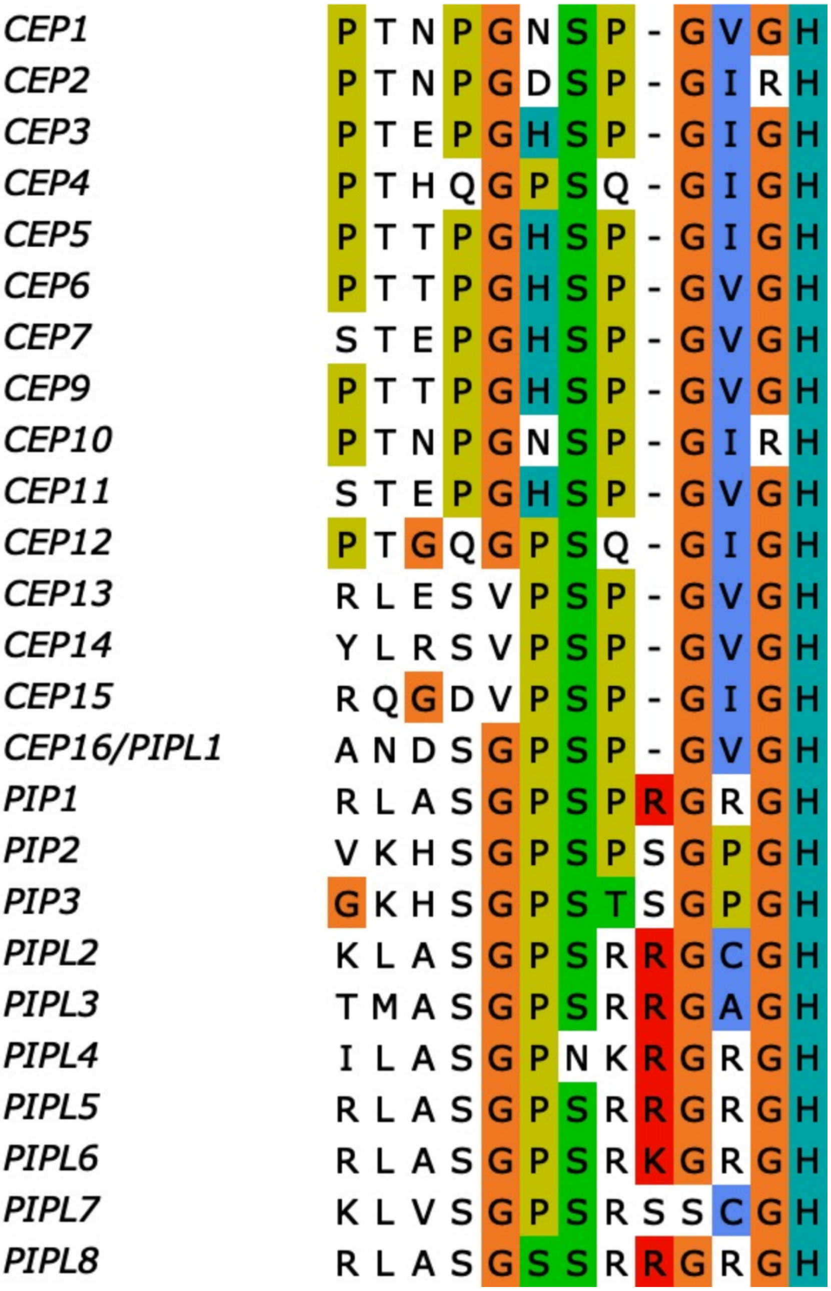


**Fig S16. Alignment of active epitopes of PIP(L) peptides.** (Related to Fig 4). Alignment of the C-terminal active epitope of CEP and PIP(L) peptides from Col-0. Alignment visualised with UGENE. Polymorphism at position 9 generates a dichotomy between PIPLs and CEPs.

Figure S17

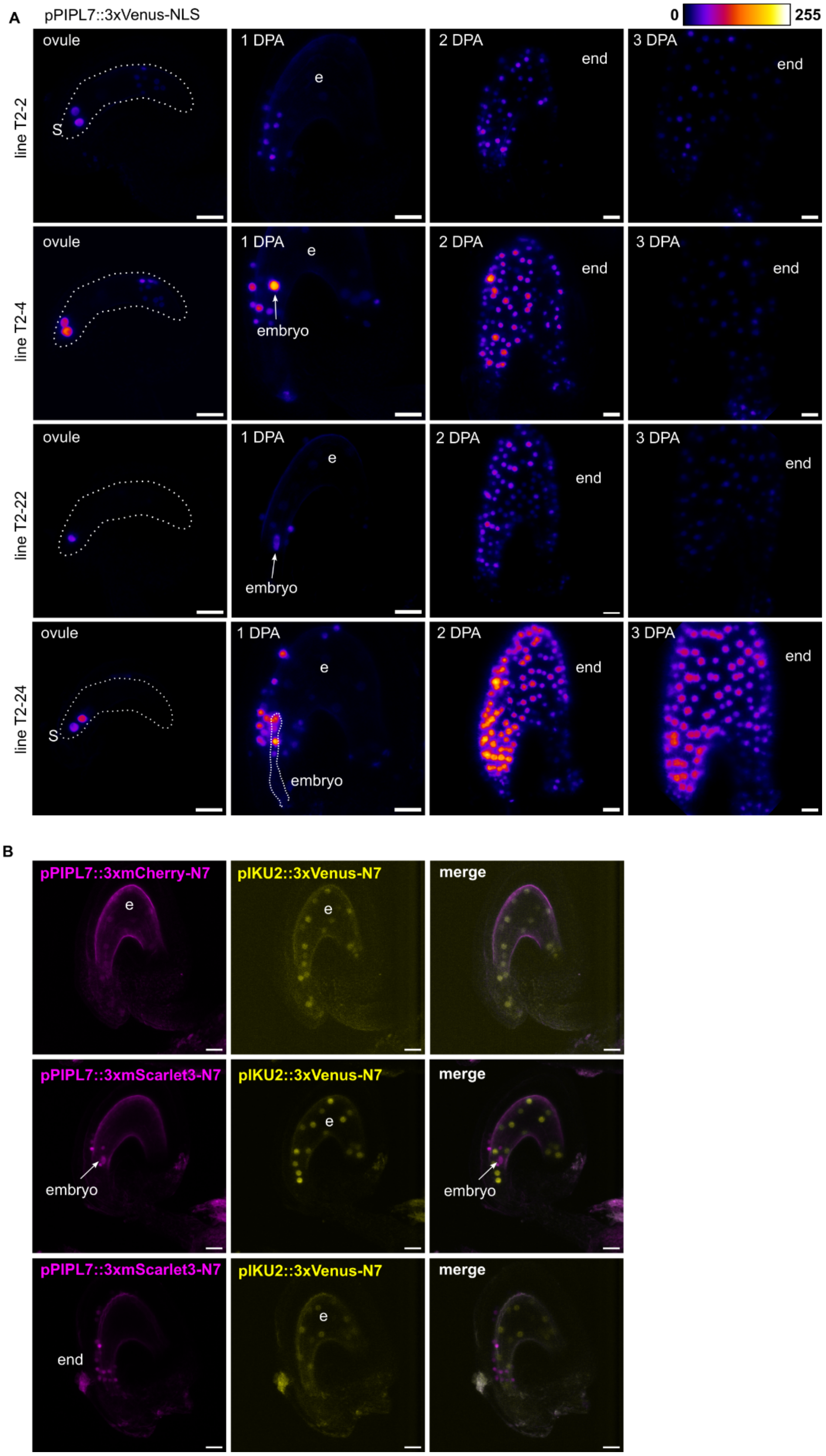
A
pPIPL7::3xVenus-NLS
0
255
line T2-2
line T2-4
line T2-22
line T2-24
ovule
1 DPA
2 DPA
3 DPA
S
e
end
embryo
B
pPIPL7::3xmCherry-N7
pIKU2::3xVenus-N7
merge
pPIPL7::3xmScarlet3-N7
e
embryo
end

**Fig S17. *PROPIPL7* expression analysis.** (Related to Fig 4). (**A**) *PIPL*7 promoter activity in Col-0 background using independent transcriptional reporter lines during early seed development; s, synergid cells; e, endosperm, end, endothelium. Representative images from n = 3-10 seeds per time point. Scale bars = 20 µm. Four independent lines are represented. (**B**) Dual transcriptional reporting of *PIPL*7 (magenta) and *IKU2* (yellow) promoter activity in Col-0 background during early seed development. e, endosperm; end, endothelium. Scale bars = 20 µm.

Figure S18

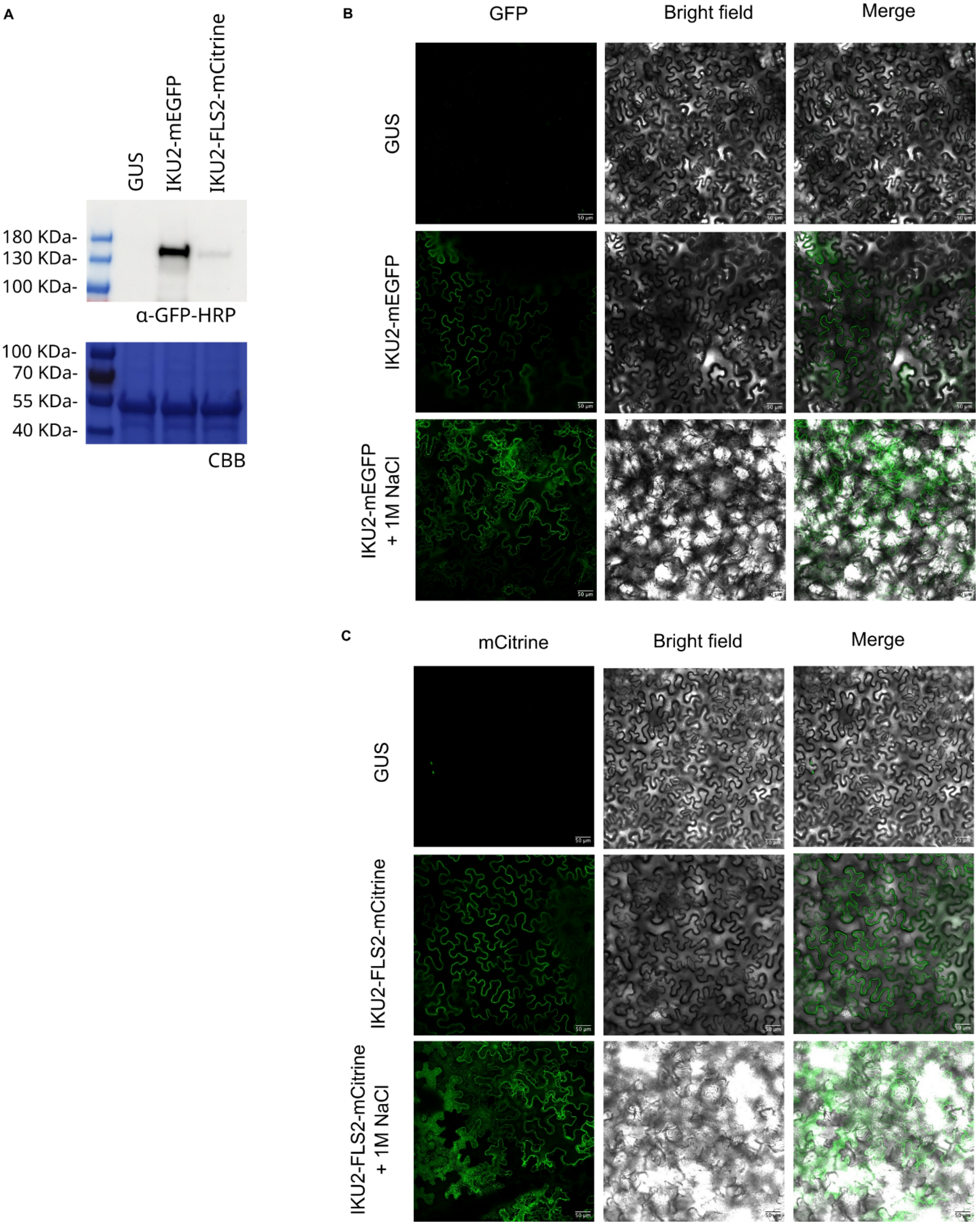
A
GUS
IKU2-mEGFP
IKU2-FLS2-mCitrine
180 KDa-
130 KDa-
100 KDa-
α-GFP-HRP
100 KDa-
70 KDa-
55 KDa-
40 KDa-
CBB
B
GFP
Bright field
Merge
GUS
IKU2-mEGFP
IKU2-mEGFP + 1M NaCl
C
mCitrine
Bright field
Merge
GUS
IKU2-FLS2-mCitrine
IKU2-FLS2-mCitrine + 1M NaCl

**Fig S18. Heterologous expression of IKU2-mEGFP and IKU2-FLS2-mCitrine in *N. benthamiana.*** (Related to Fig 4). Accumulation of IKU2-mEGFP and IKU2-FLS2-mCitrine following heterologous expression in *N. benthamiana*. (**A**) Western blot 48 h post-Agrobacterium infiltration. The western blot was probed with α-GFP-HRP as the receptor had a C-terminal mEGFP/mCitrine tags (top) and subsequently stained with CBB as a loading control (bottom). (**B-C**) Confocal microscopy (FP and Bright Field) following Agrobacterium infiltration (48 h). All confocal microscopy images in respective subpanels were recorded with the same settings and are directly comparable. The scale bar represents 50 μm. Plasmolysis was induced with 1 M NaCl for 15-30 min.

Figure S19

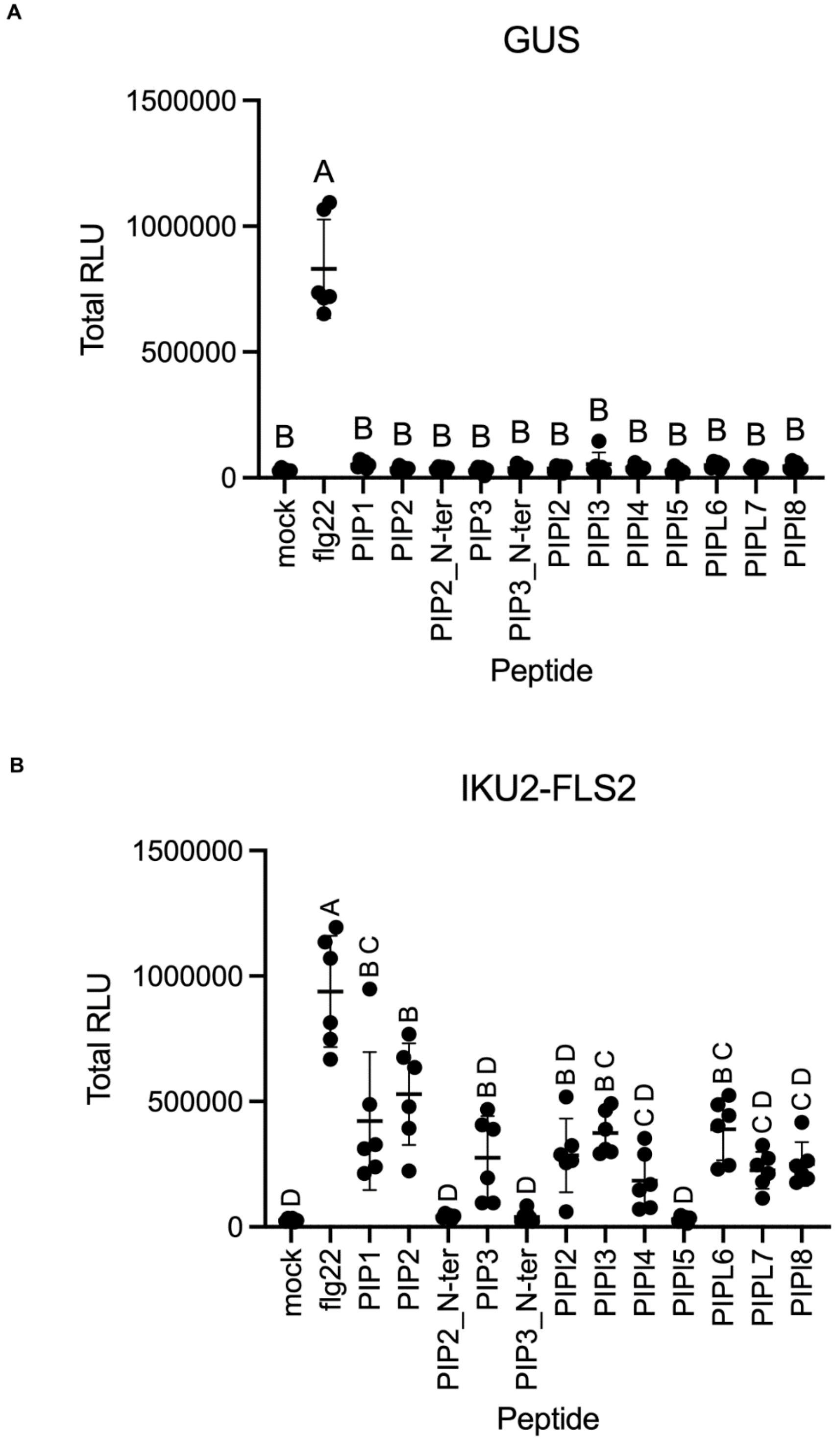


**Fig S19. Receptor-dependent elicitor-induced cytoplasmic calcium influx in *N. benthamiana*.** (Related to Fig 4). IKU2-FLS2 or GUS were transiently expressed in 35S::Aequorin transgenic *N. benthamiana* and treated with 1 µM PIPL peptides, 100 nM flg22 or mock. Assays were performed with 4-12 leaf disks per treatment and were performed independently four times with similar results. Error bars indicate S.D. Significance groupings represent One-way ANOVA giving a p-value ($<0.0001$) followed by a Tukey's multiple comparison test, the results of which are indicated as a compacted letter display.

### Figure S20

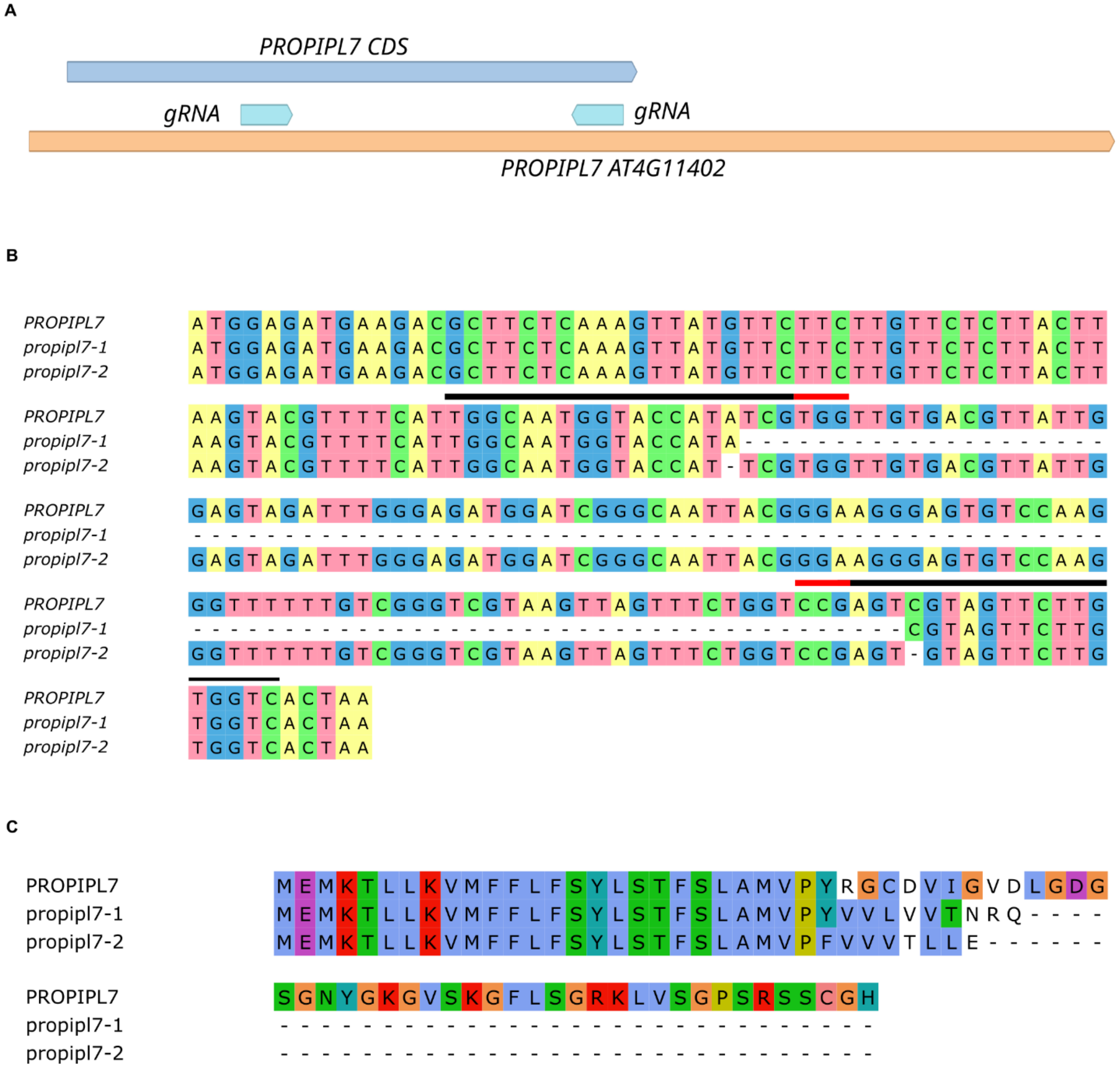


**Fig S20. Generation of *propipl7* mutants.** (Related to Fig 4). (**A**) Schematic representation of PROPIPL7 full-length transcript, coding sequence and guide RNA locations. Alignments of (**B**) DNA and (**C**) translation products of *propipl7* mutants. Black lines indicate the location of the guide RNAs, red lines indicate the PAM sites.

Figure S21

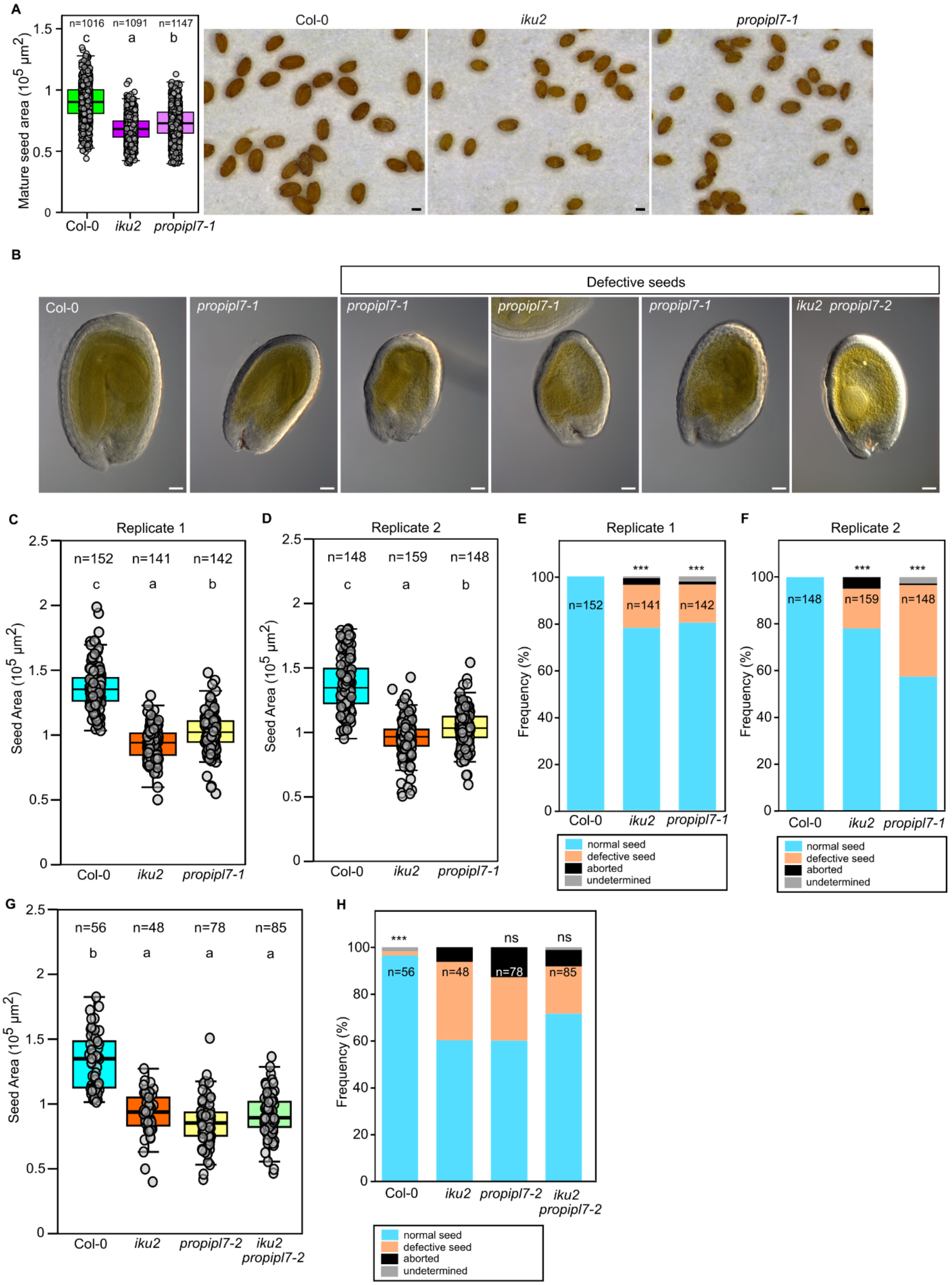
A
Mature seed area ($10^5$ µm$^2$)
n=1016 n=1091 n=1147
c a b
Col-0 iku2 propipl7-1
B
Defective seeds
Col-0 propipl7-1 propipl7-1 propipl7-1 propipl7-1 iku2 propipl7-2
C
Replicate 1
Seed Area ($10^5$ µm$^2$)
n=152 n=141 n=142
c a b
Col-0 iku2 propipl7-1
D
Replicate 2
Seed Area ($10^5$ µm$^2$)
n=148 n=159 n=148
c a b
Col-0 iku2 propipl7-1
E
Replicate 1
Frequency (%)
***
n=152 n=141 n=142
Col-0 iku2 propipl7-1
normal seed
defective seed
aborted
undetermined
F
Replicate 2
Frequency (%)
***
n=148 n=159 n=148
Col-0 iku2 propipl7-1
normal seed
defective seed
aborted
undetermined
G
Seed Area ($10^5$ µm$^2$)
n=56 n=48 n=78 n=85
b a a a
Col-0 iku2 propipl7-2 iku2 propipl7-2
H
Frequency (%)
*** ns ns
n=56 n=48 n=78 n=85
Col-0 iku2 propipl7-2 iku2 propipl7-2
normal seed
defective seed
aborted
undetermined

**Fig S21: The embryo-expressed PIPL7 peptide acts in the same genetic pathway as IKU2 to establish stable endosperm polarity.** (Related to Fig 4). (**A**) Independent biological replicate of experiment shown in Figure 4C. Left: Measurements of Col-0, *iku2* and *propipl7-1* mature seed areas from primary inflorescence stems. Total number of seeds analysed is indicated. Statistical groups established using ANOVA with Tukey's multiple comparison test; $p < 0.001$. Right: Representative images of dry seeds. Scale bars = 200 µm. (**B**) Representative Col-0, *propipl7-1* and *iku2 propipl7-2* seeds at 8 DPA, illustrating the phenotypes of defective seeds quantified in (C-F) and (G,H). Scale bars = 50 µm. (**C,D**) Measurements of Col-0, *iku2* and *propipl7-1* developing seed areas at 8 DPA from primary inflorescence stems. Two independent biological replicates are presented. Total number of seeds analysed is indicated. Statistical groups established using ANOVA with Tukey's multiple comparison test; $p < 0.001$. (**E,F**) Quantification of seed phenotypes in Col-0, *iku2* and *propipl7-1* at 8 DPA. Two independent biological replicates are presented. Data are shown as contingency bar graphs, and the total number of seeds assessed per genotype is indicated. $X^2$ test; $^{***}p < 0.001$. (**G**) Measurements of Col-0, *iku2*, *propipl7-2 and iku2 propipl7-2* developing seed areas at 8 DPA from primary inflorescence stems. (**H**) Quantification of seed phenotypes in Col-0, *iku2*, *propipl7-2 and iku2 propipl7-2* developing seed areas at 8 DPA.

Figure S22

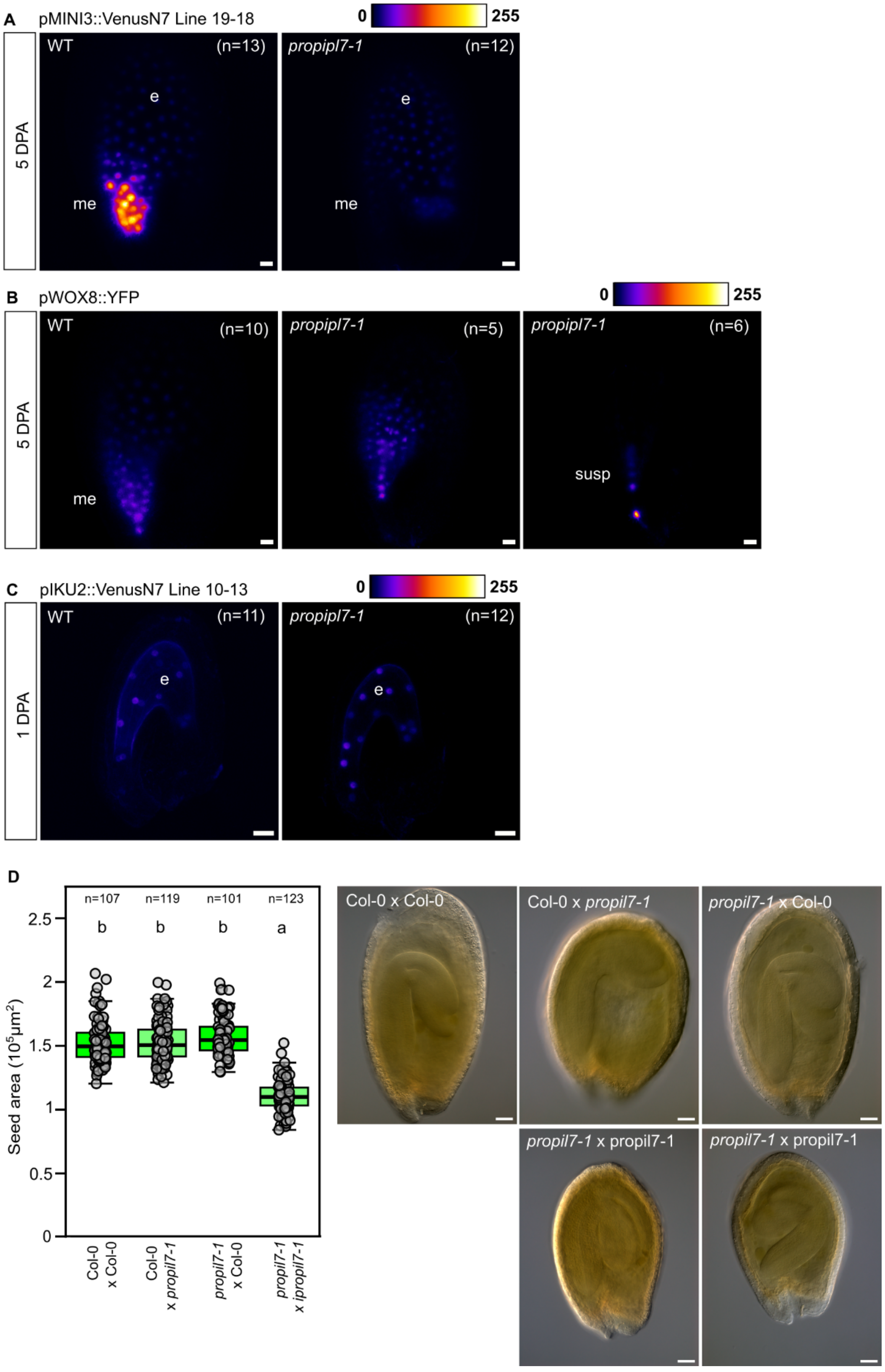
A
pMINI3::VenusN7 Line 19-18
0
255
5 DPA
WT
(n=13)
propipl7-1
(n=12)
e
me
B
pWOX8::YFP
WT
(n=10)
propipl7-1
(n=5)
propipl7-1
(n=6)
me
susp
C
pIKU2::VenusN7 Line 10-13
1 DPA
WT
(n=11)
propipl7-1
(n=12)
D
n=107
n=119
n=101
n=123
b
b
b
a
Seed area ($10^5 \mu m^2$)
2.5
2
1.5
1
0.5
0
Col-0 x Col-0
Col-0 x propil7-1
propil7-1 x Col-0
propil7-1 x ipropil7-1
Col-0 x Col-0
Col-0 x propil7-1
propil7-1 x Col-0
propil7-1 x propil7-1
propil7-1 x propil7-1

**Fig S22. The embryo-expressed PIPL7 peptide acts in the same genetic pathway as IKU2 to establish stable endosperm polarity.** (Related to Fig 4). (**A**) *MINI3* promoter activity in Col-0 and *propipl7-1* backgrounds using a transcriptional reporter line in 5 DPA seeds. (**B**) *WOX8* promoter activity in Col-0 and *propipl7-1* backgrounds using transcriptional reporter in 5 DPA seeds. (**C**) *IKU2* promoter activity in Col-0 and *propipl7-1* backgrounds using transcriptional reporter in 1 DPA seeds. (A-C) Representative images from n = 5-13 seeds. me, micropylar endosperm ;e, endosperm; susp, suspensor. Scale bars = 20 µm. (**D)** Measurement of seed areas from F1 siliques. Statistical groups established using ANOVA with Tukey's multiple comparison test; $p < 0.001$. Seeds from three independent crosses were pooled.

Figure S23

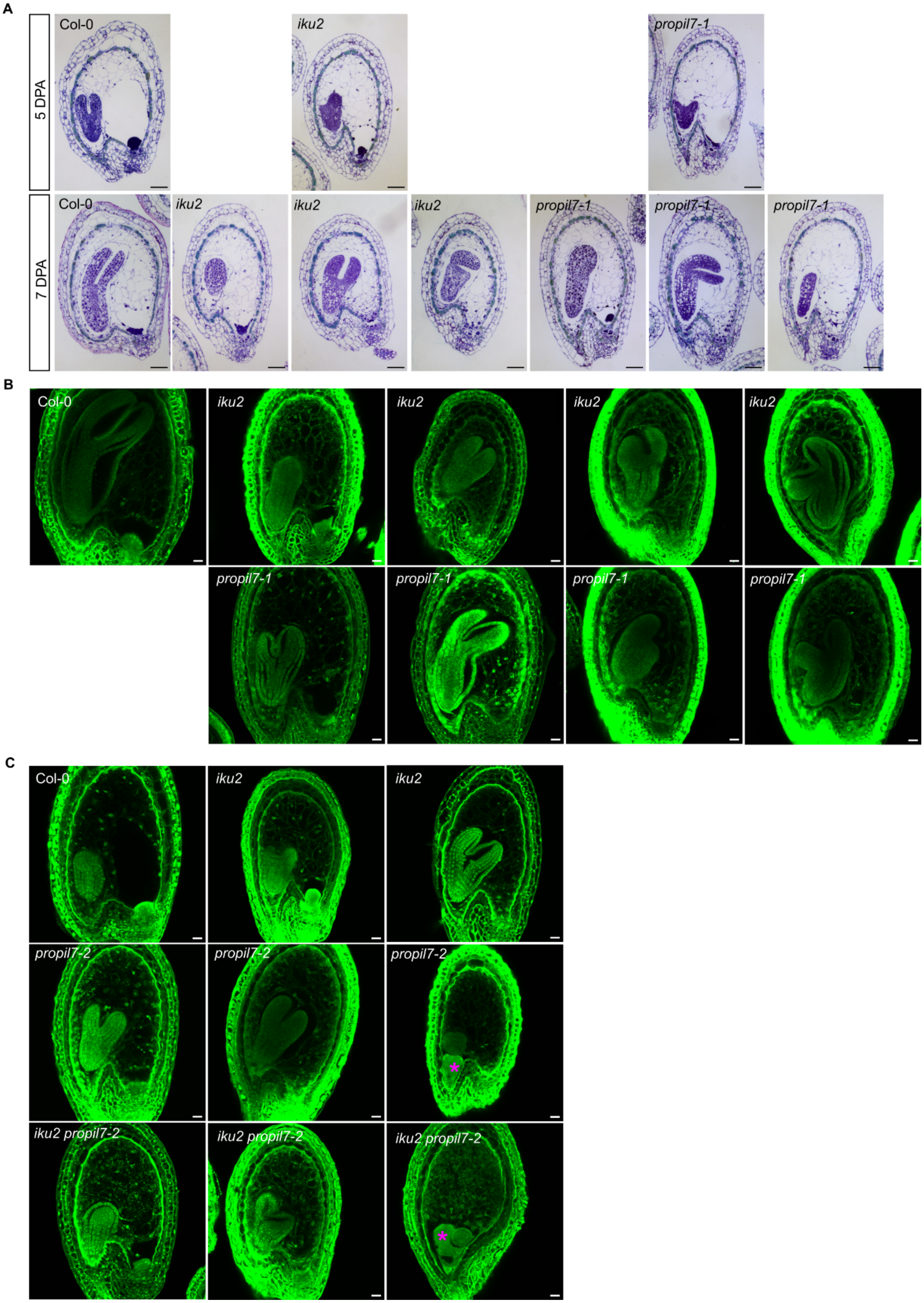

**Fig S23. The embryo-expressed PIPL7 peptide acts in the same genetic pathway as IKU2 to establish stable endosperm polarity.** (Related to Fig 4). (**A**) Representative images of median sections of Col-0, *iku2* and *propil7-1* seeds stained with toluidine blue at two stages of development. Scale bars = 50 µm. (**B**) Representative images of Col-0 and mutant seeds stained with Feulgen protocol at 7 DPA. Scale bars = 20 µm. (**C**) Representative images of Col-0, *iku2* , *propil7-2* and double mutant seeds stained with Feulgen protocol at 7 DPA. Scale bars = 20 µm. Abnormally placed endosperm cysts are indicated with magenta stars.

| Primer name | Sequence (5' -> 3') | Application |
|---|---|---|
| IKU2Delfor | TTGCTGGAGAAGCTTGTTCTAG | *iku2-2* (EMS mutant) genotyping |
| IKU2DelRev | GAACTCCATGGGAATATTCCAG | |
| SALK_050364LP | TGTGATTGAACCAATATTTGGG | *mini3-2* (SALK_050364) genotyping |
| SALK_050364RP | TCCAGCGATAACCATCGTTAG | |
| WOX8-3LP | CCCAAAATATCGTATGCATGC | *wox8-3* (SALK_014799) genotyping |
| WOX8-3RP | CCTAAACCGGAGCAGATTAGG | |
| En8130 | GCGCGTCGGTCCCCACACTTCTATAC | *wox9-1* (ZIGIA transposon) genotyping |
| t1b8-s | ATGGCTTCTTCGAATAGACACTGGC | |
| SALK LBb1.3 | ATTTTGCCGATTTCGGAAC | SALK T-DNA insertion genotyping |
| FIE-LP | TCTTTCAGTTGTGGTTCCCAC | *fie-12* (GK_362D08) genotyping |
| FIE-RP | ATGTTTCACTGAGGCCATTTG | |
| GABI-KAT LBo8474 | ATAATAACGCTGCGGACATCTACATT TT | GABI-KAT T-DNA insertion genotyping |
| EIF4A F (reference gene) | TTCGCTCTTCTCTTTGCTCTC | qPCR |
| EIF4A R (reference gene) | GAACTCATCTTGTCCCTCAAGTA | |
| At5g46630 F (reference gene) | TCAGGTGCCAATGTTCACAGC | qPCR |
| At5g46630 R (reference gene) | ACCGCTCTTCTCCCAAACCTTG | |
| MINI3 L | TGGTCAGTGATTCTTCCCAGA | qPCR |
| MINI3 R | TCTTCACCGGAAGTTTCCAC | |
| IKU2 L | GGAACTCTTGGTTACATTGCCC | qPCR |
| IKU2 R | CACCAACTCCATTAACACCACC | |
| qWOX8 Fw | CCGGCTTCAAGAATATGGTC | qPCR |
| qWOX8 Rv | GGGCTTTTGTGATGAACACG | |
| WOX9-qpF | TCGCATTCTCTCGCTACTGTCC | qPCR |
| WOX9-qpR | ACTCTTATTCGTGCGTCCGCTTG | |
| LBD35q-Fw | AGATGGAGGGACTAGATGAGGTTC | qPCR |
| LBD35q-Rv | GCGCATCAACAGGTGGTAGTTG | |
| XTH1-qRT-F | CGTACTGGTGGAATACTGGGAGTT | qPCR |
| XTH1-qRT-Rbis | TTGGAGGAACATGGAATCTAACCTTA | |

| | | |
|---|---|---|
| WOX9-S | CCAAGATGGAATCCAAAGCCAG | in situ hybridization / antisense probe |
| WOX9-T7-AS | TAATACGACTCACTATAGGGGGTCCGAAGTTGATGGGACAGTAG | |
| WOX9-T7-S | TAATACGACTCACTATAGGGCCAAGATGGAATCCAAAGCCAG | in situ hybridization / sense probe |
| WOX9-AS | GGTCCGAAGTTGATGGGACAGTAG | |
| Prom-MINI3-B4 | ggggacaactttgtatagaaaagttgAAACGGAGCATCATCGTCCAAG | MINI3 promoter cloning |
| Prom-MINI3-B1r | ggggactgctttttgtacaaacttgTTTTGACAAATCCTTAGGATGTCAAG | |
| JR1089_AT1G49800_CRISPR_1 | tgtggtctcaATTGTGGACATCATCGAGCCAAGGTTTAAGAGCTATGCTGGAA | gRNA *CEP16* |
| JR1090_AT1G49800_CRISPR_2 | tgtggtctcaATTGTATTGCGTTTGGCTCCAAGGTTTAAGAGCTATGCTGGAA | |
| JR1091_AT4G11402_CRISPR_1 | tgtggtctcaATTGTGGCAATGGTACCATATCGGTTTAAGAGCTATGCTGGAA | gRNA *PROPIPL7* |
| JR1092_AT4G11402_CRISPR_2 | tgtggtctcaATTGGACCACAAGAACTACGACTGTTTAAGAGCTATGCTGGAA | |
| ET_021_IKU2_ns_F1 | CCAAGTGGTCTCCAATGCTCCGGCTACTATTTATCG | Cloning IKU2 |
| ET_022_IKU2_ns_F2 | TTCTCGGGTCTCGGAATGGcGACCAAGGATTGCTG | |
| ET_023_IKU2_ns_R1 | CCAAGTGGTCTCCCGAaACAACTTTAGTAATCTCATCATTAGCAC | |
| ET_024_IKU2_ns_R2 | CCAATCCAAAATCAGCAATCC | |
| JR1120 | TTGGACGGTCTCGGACtTTTGACAAATGTTTCCTCTTCCCTTGgc | IKU2 ectodomain reverse primer for chimera cloning |
| JR1119 | ACAGCAGGTCTCgAGTCATCCTGATTATTCTTGGATCAGC | FLS2 cytoplasmic domain reverse primer for chimera generation |
| JR974 | ACGGATGAAGACAGCGAACCAACTTCTCGATCCTCGTTACG | FLS2 cytoplasmic domain forward primer for chimera generation |
| JR1212_mScarlet3 FWD (1) | CAGGTAGAAGACACaatgGATTCTACCGAGGCCGTG | Generation of CZLp8275 (AATG-3xmScarlet3-N7-GCTT) |
| JR1213_mScarlet3 REV (1) | CAGGTAGAAGACACTCCATAGATCCACCAGAACCACCA | Generation of CZLp8275 (AATG-3xmScarlet3-N7-GCTT) |
| JR1214_mScarlet3 FWD (2) | CAGGTAGAAGACCTTGGATTCTACCGAGGCCG | Generation of CZLp8275 (AATG-3xmScarlet3-N7-GCTT) |
| JR1215_mScarlet3 REV (2) | CAGGTAGAAGACCTAGATCCACCAGAACCACCA | Generation of CZLp8275 (AATG-3xmScarlet3-N7-GCTT) |
| JR1216_mScarlet3 FWD (3) | CAGGTAGAAGACACATCTATGGATTCTACCGAGGCCG | Generation of CZLp8275 (AATG-3xmScarlet3-N7-GCTT) |
| JR1217_mScarlet3 REV (3) | CAGGTAGAAGACACATAGATCCACCAGAACCACCA | Generation of CZLp8275 (AATG-3xmScarlet3-N7-GCTT) |
| JR1218_NLS_Fw | CAGGTAGAAGACTCCTATCGCTGCAGCGGCCGAA | Generation of CZLp8275 (AATG-3xmScarlet3-N7-GCTT) |
| JR1219_NLS_Rv | CAGGTAGAAGACTCaagcTTACTCTTCTTCTTGATCAGCTTCTGTG | Generation of CZLp8275 (AATG-3xmScarlet3-N7-GCTT) |

| Plasmid identifier | Plasmid description |
|---|---|
| CZLp6841 | LP CRISPR Cas9 construct targeting AT4G11402 and AT1G49800 |
| CZLp6966 | 35S::IKU2-FLS2-mCitrine |
| CZLp4332 | 35S::IKU2-mEGFP |
| CZLp7748 | *GGAG-CP selection casette-AATG 3xVenus-N7 CDS* |
| CZLp7320 | GGAG-*pPROPIPL7*-AATG |
| CZLp8275 | AATG-3xmScarlet3-N7-GCTT |
| CZLp8271 | pPROPIPL7::3xmScarlet3-N7 |
| CZLp8270 | pPROPIPL7::3xmCherry-N7 |
| ACpH7-LR2 | pMINI3::3xVenus-N7 |

| Peptide name | Peptide sequence |
|---|---|
| >flg22 | QRLSTGSRINSAKDDAAGLQIA |
| >PIP1 | RLASG{Hyp}SPRGPGH |
| >PIP2 | VKHSG{Hyp}SPSGPGH |
| >PIP2_N-ter | IKDSG{Hyp}S{Hyp}GEGH |
| >PIP3 | GKHSG{Hyp}STSGPGH |
| >PIP3_N-ter | IKESG{Hyp}SSGGEGH |
| >PIPL2 | KLASG{Hyp}SRRGCGH |
| >PIPL3 | TMASG{Hyp}SRRGAGH |
| >PIPL4 | ILASG{Hyp}NKRGRGH |
| >PIPL5 | RLASG{Hyp}SRRGRGH |
| >PIPL6 | RLASG{Hyp}SRKGRGH |
| >PIPL7 | KLVSG{Hyp}SRSSCGH |
| >PIPL8 | RLASGSSRRGRGH |

N.B. All peptides were synthesised by Genscript to a purity >85%

#### Table S1. Primers, plasmids and peptides generated in this study.
